# Weight and organ specific immune cell profiling of Sleeve Gastrectomy

**DOI:** 10.1101/2020.06.28.176628

**Authors:** David A. Harris, Renuka Subramaniam, Todd Brenner, Ali Tavakkoli, Eric G. Sheu

## Abstract

Sleeve gastrectomy (SG) has profound, immediate weight-loss independent effects on obesity related diabetes (T2D). Our prior studies have shown that immunologic remodeling may play a part in this metabolic improvement. However, to date, little is known about how the major immune cell populations change following SG. Using mass cytometry with time of flight analysis (CyTOF) we aimed to broadly explore the organ-specific immune cell repertoire induced by SG. Surgery was performed on obese, insulin resistant and lean mice in order to understand surgery-specific phenotypes. We identified a shift within the splenic B cell compartment with a reduction in follicular and an increase in innate-like B cell subsets in SG animals. There was a concomitant increase in multiple circulating immunoglobulin classes. Further, SG animals had a conserved increase in splenic neutrophils and a tendency toward M2 macrophage polarization. Others have shown that these, weight-loss independent, surgery-specific changes are linked to improved glucose metabolism and thus, may be a major contributor to post SG physiology. Characterizing the complex immune milieu following SG is an important step toward understanding the physiology of SG and the potential therapies therein.

## Introduction

Sleeve gastrectomy (SG) and other forms of bariatric surgery are the most successful therapies for obesity-related type 2 diabetes (T2D). While patients experience significant benefit from weight loss, our group and others have shown that improved glycemic control often occurs independent of weight loss following SG.(1–4)

Despite a large effort to understand the scientific underpinnings of SG, there has yet to be a unifying theory on SG-induced T2D remission. To better understand the immediate post-SG physiology, we developed a novel model of sleeve gastrectomy in lean mice. In this model, SG does not lead to sustained changes in weight or body composition. Metabolic caging, PET/CT imaging, and RNAseq revealed increases in systemic and local visceral adipose tissue glucose utilization and surprisingly, immunologic remodeling as a potential driver of this process.(5)

Obesity induces a chronic, low-grade immune dysregulation in the form of increased circulating pro-inflammatory cytokines and increased Th1 T cells, CD8+ T cells, macrophages, neutrophils, and dendritic cells (DC). These directly contribute to obesity-related insulin resistance.(6–11) In mice with diet induced obesity (DIO), accumulation of M1-polarized macrophages within white adipose depots contributes to chronic inflammation.(6)

Similarly, accumulation of B2 cells and IgG2c within visceral adipose tissue (VAT) in DIO mice is sufficient to confer insulin resistance. Transfer of either to lean B^null^ mice leads to similar impaired glucose metabolism.(8) B cells also promote insulin resistance through production of proinflammatory cytokines and direct T-cell interactions.(12) By contrast, alternative B cell programming mitigates obesity-induced T2D through production of IL-10 and endogenous immunoglobulin while affecting adipose tissue macrophage polarization.(13–17) Together, these data indicate that B cells both contribute to, and ameliorate, obesity and T2D through antibody production and regulation of variable cytokine production.

Bariatric surgery affects obesity-induced inflammation through upregulation of adiponectin and IL-10, reduction in pro-inflammatory cytokines, and induction of a reparative T cell phenotype.(18) We showed that SG in rats reduced jejunal IL-17 and IL-23 as well as distal jejunal INFγ, which correlated with weight loss and improved glucose handling.(19) Additionally, in a longitudinal human study of SG-induced circulating immunologic phenotype, we identified a reduction in the neutrophil to lymphocyte ratio, circulating IL-6, and circulating CRP levels across subjects (unpublished). Others showed that SG leads changes in macrophage, T cell, and DC function within VAT compared to weight-matched controls and that surgery can induce alternative B cell programming.(20–22)

While these studies offer important insight into the post-SG immune state, it is hard to establish the relative importance of these findings against the backdrop of a globally changing immune system induced by both the immediate SG-induced physiology and weight loss. To date, there are no comprehensive studies evaluating local and systemic immune cell phenotypes following SG. Thus, we employed a novel multiplex platform, cytometry with time of flight analysis (CyTOF), to characterize and quantify changes across the organ-specific immune cell fractions.(23) We leveraged lean and DIO models of SG to separate the surgery-specific and weight-loss specific immune phenotype in SG animals.

## Methods

### Animals

11-week-old, male, lean or insulin resistant/obese (DIO) C57Bl/6J mice reared on standard chow (5% calories from fat; Pico5053, Lab diet, St. Louis, MO) or high fat diet (60% calories from fat; RD12492, Research diets, New Brunswick, NJ), respectively, were purchased from Jackson Laboratory (Bar Harbor, ME). They were housed in a pathogen-free, climate-controlled environment with a 12-hour light/dark cycle and maintained on their respective diets. Figure 1. outlines the experimental design. After 1-week acclimation, mice underwent operations as outlined below. Animals were cared for according to the American Association for Laboratory Animal Science. Procedures approved by our Institutional Animal Care and Use Committee.

**Figure 1:**
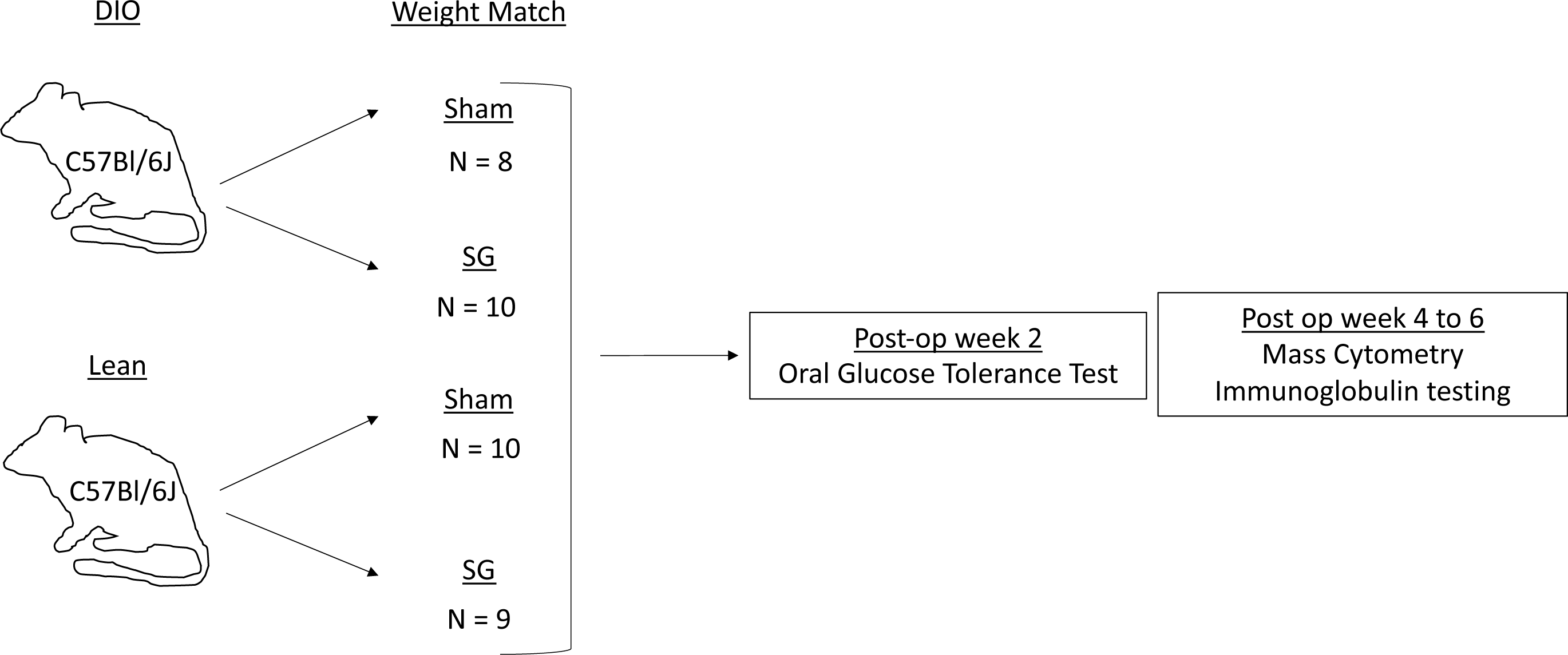
Two experiments were completed for both DIO and Lean groups. DIO experiment 1: n = 4, 6 for sham and SG, respectively. Cytometry completed on jejunum, liver, spleen. DIO Experiment 2: n = 4, 4. Cytometry completed on Jejunum, ileum, cecum, and spleen. Lean experiment 1: n = 5, 4. Cytometry completed on jejunum, ileum, liver, and spleen. Lean experiment 2: n = 5, 5. Cytometry completed on jejunum, ileum, cecum, and spleen.

### SG and Sham operations

Lean and DIO mice were weight matched within their groups and underwent either SG or Sham operations. SG consisted of short gastric vessel ligation and resection of 80% of the stomach including the entirety of the non-glandular portion. Sham operation consisted of a similar laparotomy, vessel ligation, and manipulation of the stomach along the resection equivalent. Mice were then maintained on Recovery Gel Diet (Clear H_2_O, Westbrook, ME) for 6 days and then restarted on their original diet formulation. Four separate experiments were performed – two lean (n = 5, 4 and n = 5, 5 for sham and SG, respectively) and two DIO (n = 4, 6 and 5, 4 for sham and SG, respectively).

### Oral glucose tolerance tests (OGTT)

Performed at 2 weeks post-operatively. After a 4 hour fast (8 am to noon), mice were orally gavaged with 2mg/g of oral D-Glucose (Sigma-Aldrich, St. Louis, MO). Serum glucose was measured from the tail vein with a OneTouch Glucometer (Life Technologies, San Diego, CA).

### Preparation of single cell suspension

Animals were sacrificed 4 to 6 weeks post-operatively. Blood was collected via cardiac puncture. The right ventricle perfused with 30mL of Hank’s Balanced Salt Solution (HBSS) containing 5% fetal bovine serum (FBS), 10mM HEPES, and 0.2% heparin (Life Technologies). The liver, spleen, proximal jejunum, distal ileum, and cecum were harvested. Single-cell suspensions were prepared using Liberase TL (Roche, Penzberg, Germany; 200 ng/mL) and DNAse I (Sigma-Aldrich; 0.15 mg/mL). Lysates were strained through a 70uM strainer (Celltreat, Pepperell, MA). Red blood cell lysis was performed in ACK lysis buffer (Life Technologies). This was not performed on liver samples due to high cell death rates. Cells were counted using a hemocytometer and 0.4% trypan blue (Life Technologies) and aliquoted into 96-well polypropylene plates at a density of 1 million cells per well.

### Cell staining

Cells were suspended in cell staining buffer (CSB - CyTOF-grade PBS, 0.625% BSA, and 25mg sodium azide; all from Sigma). 100μM cisplatin was used for viability staining (Fluidigm, San Francisco, CA). Cells were blocked with mouse Fc block (BioLegend, San Diego, CA). Staining for extracellular markers was performed for 30 min at room temperature. Cells were fixed and permeabilized using permeabilization buffer (eBioscience, San Diego, CA) and stained for the intracellular markers for 30 min at room temperature. Cells were fixed in 1.6% paraformaldehyde (Thermo Fisher Scientific, Waltham, MA), washed, and stained with iMaxPar Intercalator-Ir (Fluidigm). They were resuspended in water with EQ Four Element Calibration beads (Fluidigm) at a density of 100,000 to 1,000,000 cells/mL and strained through 35μm filter (BD Biosciences, Bedford, MA).

### CyTOF Acquisition Methods

Samples were analyzed at the Dana Farber Cancer Institute, Longwood Medical Area CyTOF Core. The Helios (Fluidigm) mass cytometer was auto-tuned with CyTOF tuning solution (Fluidigm) with mass resolution>400, 159Tb dual counts >600,000, detector voltage <-1300V, dual slopes =0.03±0.003, R2>0.8, oxide ratio (M1/M2)<0.03, and percent RSD for tuning solution elements <3%. A bead sensitivity test was performed using EQ(tm) Four Element Calibration Beads. The percent singlet was >95%, mean intensity of 153Eu was >1500, cv of 153Eu <15%, and the cv of 156Gd <100%. Samples were normalized post-acquisition using CyTOF software (Fluidigm, CyTOF version 6.7.1014) by using the known intensities of the Calibration Beads in each sample such that bead intensities matched the bead passport (EQ-P13H2302_ver2).

### Panel Design

A 29-antibody panel allowed for simultaneous evaluation of major innate and adaptive immune populations – neutrophil, macrophage, dendritic cell, natural killer (NK) cell, innate lymphoid cell (ILC), T cell, and B cell. Antibodies were conjugated and validated by the Harvard Medical Area CyTOF Antibody and Resource Core. We had two separate iterations of our panel. In the second shown in Supplemental figure 1, CD80 and IgD were added and absent in the first lean and DIO experiments.

### Data Analysis

Cytobank software suite was used for analysis (Cytobank, Mountain View, CA). High-dimensional cytometry data analysis was performed using the visualization of t-distributed stochastic neighbor embedding (viSNE) algorithm.(24) Traditional Phenotyping was also performed using the following gating strategies of the CD45^+^ leukocyte lineage (Supp figure 1). T cells: CD3^+^ and positivity for either CD4 or CD8 with subdivision based on Tbet, RORγt, GATA3, and FoxP3. NKT cell: CD3^+^ and subdivision via Nk1.1 and NKp46 (Supp figure 2). B cells: CD19^+^CD11B^+/neg^ and subdivision via CD21, CD23, IgM, and IgD.(8,16,25,26) NK cell: CD3^neg^ and subdivision via Nk1.1 and NKp46. A negative lineage gate (CD3^neg^CD5^neg^NK1.1^neg^NKp46^neg^CD19^neg^CD11b^neg^CD11c ^neg^) was used to identify ILC populations, which were divided into ILC1, ILC2, and ILC3 by the expression of Tbet, GATA3, and RORyt, respectively (Supp figure 3). Neutrophils: CD11B^+^Ly6G^+^. Macrophages: CD11B^+^F4/80^+^ cells with subdivision via CD80, MHCII, Ly6C, and Arginase-1 (ARG1).(27–29) Dendritic cells: CD11C^+^MHCII^+^Ly6G^neg^ (Supp figure 4).(30) Unpaired, two-tailed Student’s T-test (p = 0.05) were used to compare populations as a percentage of total CD45^+^ leukocytes in SG and Sham animals. Assessment of Arg1 staining was made by comparing the mean intensity of antibody staining (unpaired, two-tailed Student’s T-test, p = 0.05).

**Figure 2:**
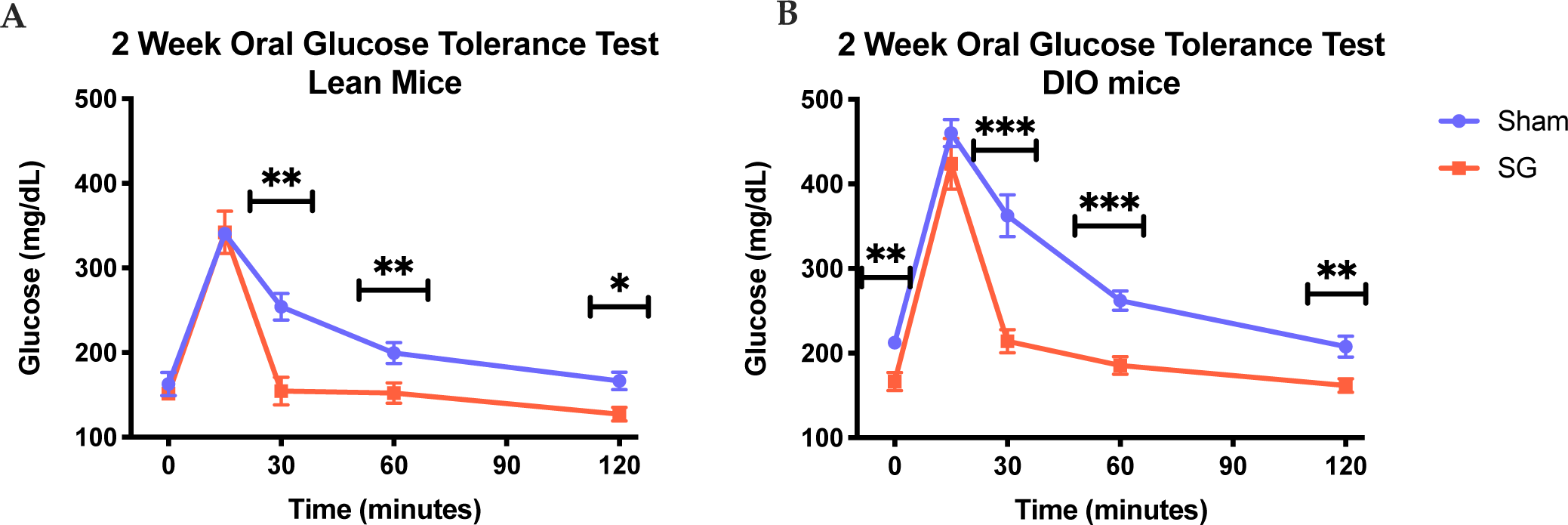
Oral glucose tolerance test in Lean (A, n=5,5) and DIO (B, n=10,10) mice 2 weeks following sham or SG. Error bars represent SEM. Student’s T-test. *p<0.05, **p<0.01, ***p<0.001

**Figure 3:**
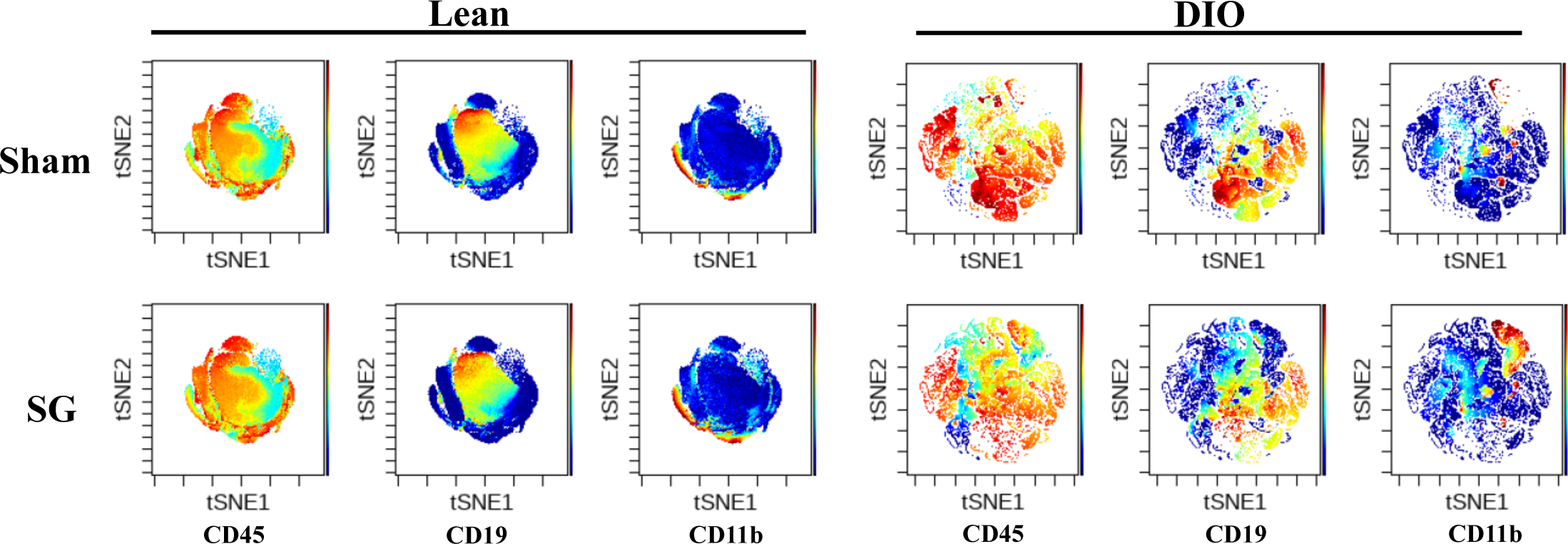
Multivariate viSNE analysis demonstrates qualitative alterations in the splenic immune milieu after SG in both lean and obese animals. Representative plots are colored by signal intensity of CD45, CD19, and CD11b respectively.

**Figure 4.**
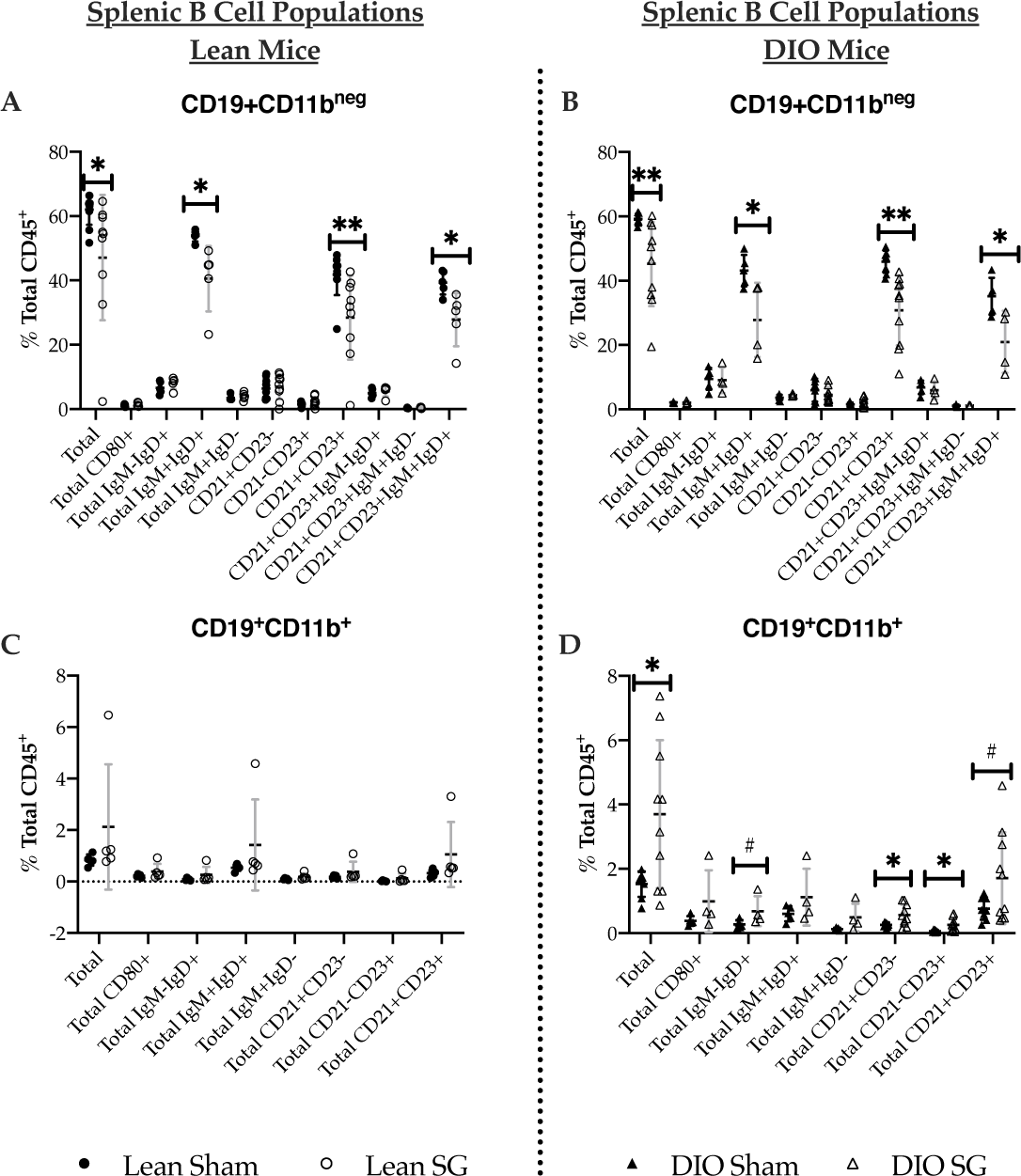
CD11b negative and positive B cell populations in Lean (A and C, respectively) and DIO (B and D, respectively) sham and SG mice. Significance was determined by Student’s T test. #p<0.10,*p<0.05, **p<0.01. Means with standard deviation superimposed.

## Quantitation of immunoglobulins

Sera were collected from Lean and DIO mice (n=6 per group) 4 to 6 weeks following surgery. Total IgM, IgG, and IgG subtypes were measured by ELISA (Thermo Fisher Scientific).

## Data and Resource Availability

The datasets generated during the current study are available from the corresponding author upon reasonable request. The antibodies used during the current study are available from the Harvard Medical Area CyTOF Antibody and Resource Core.

## Results

### SG leads to improved glucose handling in DIO and Lean mice

We have previously shown that lean mice have altered glucose handling following SG independent of weight loss and lasting changes body composition or caloric intake.(5) However, unlike lean mice, SG in DIO mice leads to improved glucose handling with concomitant weight loss.(31) Thus, leveraging these two models, we attempted to separate weight-independent and dependent effects of SG.

In the present study, we again demonstrate that Lean and DIO mice have a conserved response to oral glucose challenge at 2 weeks post-procedure. Fasting glucose levels were not different between Lean sham and SG animals (115-200mg/dL) but DIO SG animals had a lower fasting glucose levels than DIO Shams (166±36 vs 212±23mg/dL, p=0.002, Figure 2). However, Lean and DIO SG animals both have a rapid reduction in serum glucose levels toward baseline between 15- and 30-minute compared to respective sham animals.

We have also previously shown that SG in DIO and Lean animals leads to improved insulin sensitivity as measured by insulin tolerance testing and increased levels of the glucoregulatory hormone, GLP-1 despite equal caloric intake between sham and SG animals in each respective cohort.(5,31)

### viSNE reveals perturbations in B cell and CD11B^+^ subsets following SG in Lean and DIO mice

CyTOF data were visualized on two axes using the viSNE algorithm-generated variables tSNE1 and tSNE2. Figure 3 shows representative plots of total splenic leukocyte populations from Lean and DIO operative groups. In Lean and DIO SG animals there is a reduction in total CD19^+^ and an increase in total CD11b^+^ leukocytes compared to their sham counterparts. The latter includes myeloid cell and innate-like lymphocyte populations. The dominant CD11b^+^ cluster stained for neutrophil, macrophage, and dendritic cell markers such as F4/80, MHCII, CD11c, and Ly6G. Interestingly there was an increase in CD11b^+^ within the CD19^+^ cluster in DIO SG animals. There were no clear differences in viSNE analysis of the liver, ileum, cecum, and jejunum.

### B cell populations in DIO and Lean mice

Given the qualitative evidence supporting a change in the splenic B-cell compartment following SG in both lean and DIO mice, we next performed gated analysis to assess specific B cell population changes. B cells were first categorized as being CD19^+^ and CD11b^neg^ or CD11b^+^. CD11b is important for B-cell migration, tissue infiltration, and B cell receptor modulation.(32–Subsets were then created based on expression of CD21, CD23, IgM, and IgD in order to delineate marginal zone (MZ) from follicular and mature from immature B cells.(8)

In lean animals, SG was associated with a 15% reduction in total splenic CD19^+^CD11b^neg^ B cells (p=0.031) with a concomitant 14%, 13%, and 12% reduction in IgM^+^IgD^+^ (p=0.020), CD21^+^CD23^+^ follicular (p=0.010), and mature CD21^+^CD23^+^IgM^+^IgD^+^ follicular cells (p=0.022), respectively (figure 4a). There were no differences in CD21^+^CD23^-^ or CD23^+^CD21^-^ subsets in lean SG mice. There were also no changes in jejunal, ileal, cecal, or hepatic CD19^+^CD11b^neg^ B cell subsets. Finally, there were no difference in CD19^+^CD11b^+^ B cell populations in ileal, cecal, splenic, or hepatic tissues. In jejunal specimen, SG was associated with a small reduction in CD19^+^11b^+^IgM^+^IgD^+^ B cells (p=0.010; Supplemental table 5)

Similarly, in DIO animals, SG was associated with a 14% reduction in total splenic CD19^+^CD11b^neg^ B cells (p=0.009) with a concomitant 15%, 15%, and 14% reduction in IgM^+^IgD^+^ (p=0.030), CD21^+^CD23^+^ follicular (p=0.002), and mature CD21^+^CD23^+^IgM^+^IgD^+^ follicular subsets (p=0.028), respectively (figure 4b). Additionally, SG animals had a 2.0-fold reduction and 2-fold increase in hepatic CD19^+^CD11b^neg^CD21^+^CD23^neg^ (p=0.017) and CD19^+^CD11b^neg^CD21^neg^CD23^+^ B cells (p=0.038), respectively. There were no differences in jejunal, ileal, or cecal CD19^+^CD11b^neg^ B cells populations (supplemental table 10).

Unexpectedly, DIO SG mice had an increase in multiple splenic CD19^+^CD11b^+^ populations. CD11b^+^ B cells are Innate-like B cells found within the splenic marginal zone (MZ) and peritoneal cavity.(32) There was a increase in total CD19^+^CD11b^+^ (2.5-fold; p=0.019), CD21^+^CD23^neg^ (2.2-fold; p=0.022), and CD21^neg^CD23^+^ (4.3-fold; p=0.015) innate-like B cells (figure 4d). There was also a 2-fold increase in jejunal CD19^+^CD11b^+^IgM^+^IgD^neg^ B cells (supplemental table 11). While there were no significantly different changes in these populations with Lean SG mice, there were similar increases in sub-populations as identified in DIO SG mice (figure 4c; supplemental table 5)

### Myeloid cells

While there was no difference in total macrophage or macrophage subsets in DIO SG and sham mice (supplementary table 12), lean SG mice had a 2.9-fold increase in splenic CD11c^+^MHCII^neg^ macrophages (p=0.029) and an increase in multiple ileal macrophage subsets compared to lean shams. There were no differences in Ly6C expression total macrophage populations from Lean or DIO mice within jejunal, cecal, splenic, and hepatic tissue. Lean SG animals had a 2.7-fold, 3-fold, and 4.2-fold increase in total Ly6C^neg^ (p=0.011), CD11c^+^MHC^lo^ (p=0.016), CD11c^neg^MHCII^neg^ (p=0.006) macrophages, respectively (supplementary table 6)

We next examined macrophage polarization via CD80 (classically activated, pro-inflammatory, M1) and Arg1 (alternatively activated, reparative, M2) expression.(28,29) In Lean SG mice, there was a 3.1-fold, 2.1-fold, and 3.6-fold increase Arg1 expression in total splenic F480^+^ (p=0.011), CD11c^+^MHCII^+^ (p=0.020), and CD11c^+^MHCII^lo^ (p=0.037) macrophages (figure 5a). Additionally, there was a 4.5-fold and 6.8-fold increase in Arg1 expression in total and CD11c^neg^MHCII^+^ ileal macrophages and a 1.1-fold and 1.2-fold increase in splenic total F480^+^ (p=0.003) and CD11C^+^MHCII^+^CD80^+^ (0.034) macrophages (supplemental table 7). In DIO SG mice, there was a 1.8-fold and 1.4-fold increase in Arg1 expression in splenic CD11c^+^MHCII^+^ (p=0.046) and CD11c^neg^MHCII^neg^ (p=0.028) macrophages, respectively (figure 5b). There were no changes in the number of CD80^+^ macrophages across any organ tested (supplemental table 13).

**Figure 5.**
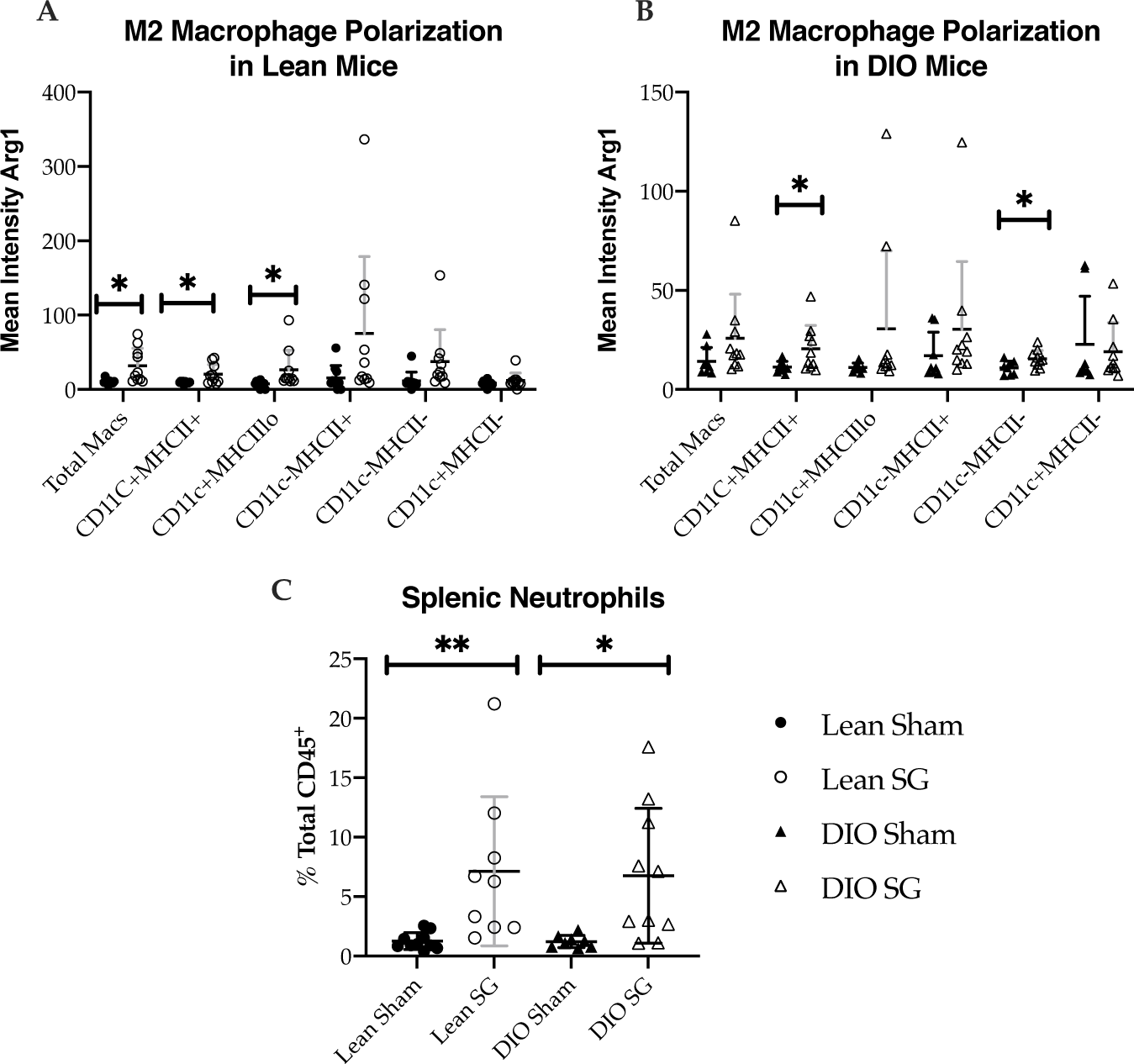
M2 macrophage polarization as measured by mean intensity staining of Arg1 in Lean (A) and DIO (B) Sham and SG mice. (C) Quantification of splenic neutrophils. Significance was determined by Student’s T test. *p<0.05, **p<0.01. Means with standard deviation superimposed.

Interestingly, there was a conserved 5.5-fold (p=0.009) and 5.7-fold increase in total Ly6G^+^CD11B^+^ neutrophils in splenic specimen from both lean and DIO SG mice, respectively (figure 5c). Both also saw an increase in gut neutrophil populations with a 3-fold increase in ileal neutrophils in Lean SG mice (p=0.014) and 4-fold increase in cecal neutrophils in DIO SG mice (p=0.013). There were no other conserved changes in myeloid populations between DIO and Lean SG.

### Additional Immune subsets in Lean and DIO Sham and SG animals

No clear patterns of expression were identified in jejunal, ileal, cecal, hepatic, and splenic T cell, NK, NKT cell, and ILC subsets. Lean SG mice had a 1.4-fold reduction in both total T CD3^+^ cells (p=0.009) and CD4^+^ T cells (p=0.013; supplemental table 2). DIO SG mice had a 1.6-fold reduction in splenic CD8^+^ T cells (supplemental table 8). Lean SG mice had a 2.0-fold and 2.7-fold increase in splenic CD3^neg^NK1.1^+^ (p=0.016) and cecal CD3^neg^Nk1.1^+^NKp46^+^ (p=0.039) NK cells, respectively (supplemental table 3). DIO SG mice had a 1.4-fold decrease in splenic CD3^neg^Nk1.1^+^NKp46^+^ (p=0.006) NK cells (supplemental table 9).

### SG is associated with increased immunoglobulins

Given the significant but opposing changes in CD11b^+^ and CD11b^neg^ B cell subsets, we next assessed for variance in the circulating immunoglobulin profile between SG and Sham animals in Lean and DIO states. Interestingly, despite reduced numbers in the vast majority of B cell subsets in SG animals, there was an increase in total serum IgM and IgG in Lean (figure 6a: 1.5-fold and 3.6-fold, respectively) and DIO SG animals (figure 6b: 1.5 fold and 5.5-fold, respectively) as compared to Shams. Further exploration of IgG subtypes showed that SG was associated with a 4.4-fold increase in IgG2b and 2.2-fold increase in IgG3 with a trend toward increased IgG1 in Lean mice (figure 6c) and a 1.7-fold increase in IgG2b and 3.8-fold increase in IgG2c with a trend toward increased IgG1 and IgG3 in DIO mice (figure 6d)

**Figure 6.**
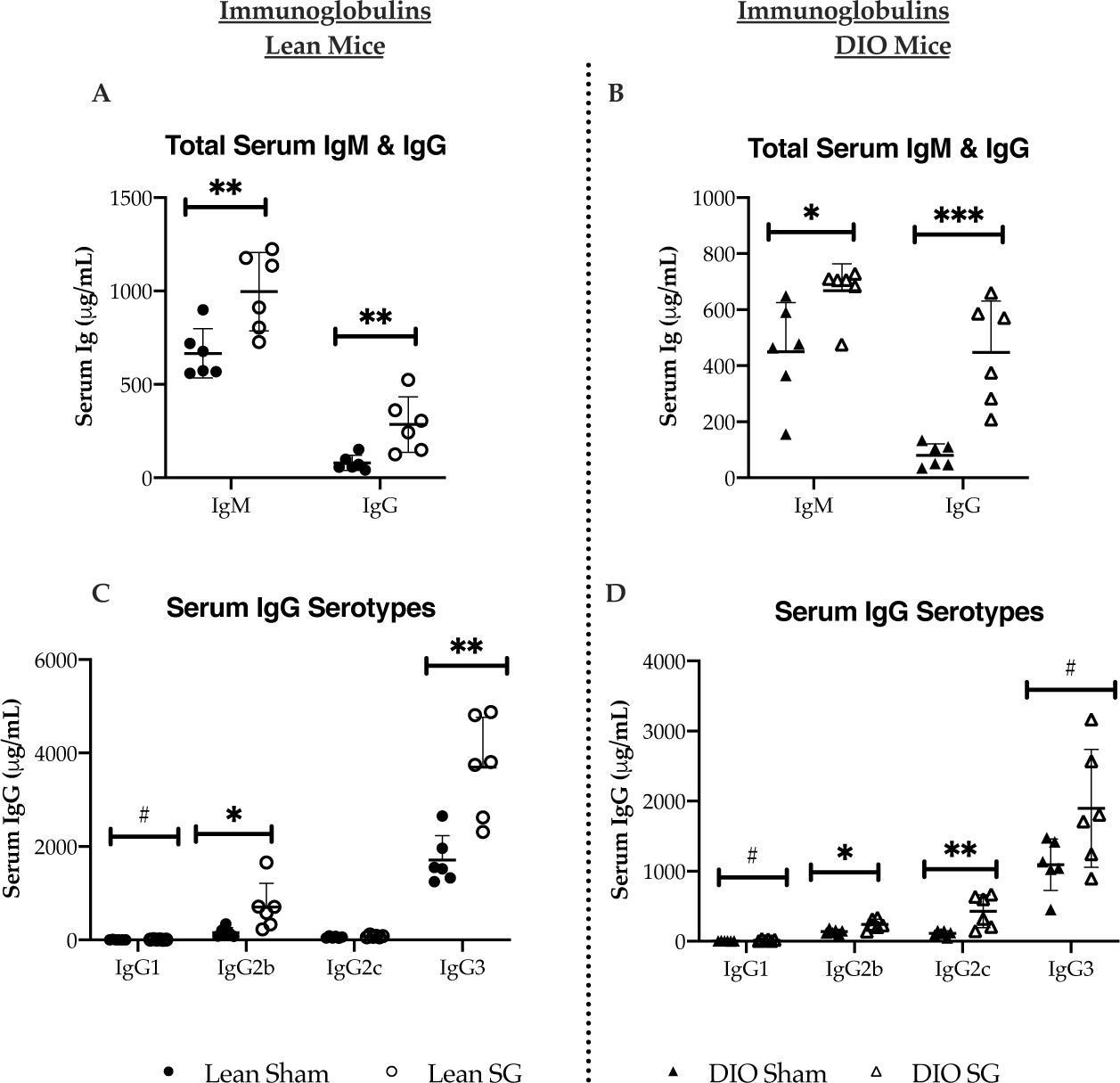
Total immunoglobulin and IgG serotypes in Lean (A and C, respectively) and DIO (B and D, respectively) sham and SG mice. Significance was determined by Student’s T test. #p<0.10,*p<0.05, **p<0.01, ***p<0.001. Means standard deviations superimposed.

## Discussion

SG has profound weight-loss independent and dependent effects on glucose metabolism. In our prior work, we developed a novel model of SG in lean C57Bl/6J mice reared on normal chow to understand the immediate effects of surgery without the influence of diet and weight-loss. Utilizing indirect calorimetry and PET-CT imaging, we found that SG induces a shift toward preferential glucose utilization, in part, driven by increased glucose uptake within the white adipose tissue depots. Gene expression analysis of the visceral adipose tissue revealed a surprising upregulation in immune-centric genes, pathways, and transcription factors.(5) In the present study, we follow-up these results by characterizing weight dependent and independent SG-induced changes in immune cell populations by way of the CyTOF multiplex immune platform.(23)

We observe several major cellular immune changes conserved across both Lean and DIO SG models. First, SG induced changes in B lymphocyte subpopulations. Including, a global reduction in CD19^+^CD11b^neg^ B cells and related follicular B cell subsets, increased innate-like, CD11B^+^ B cell populations, and increased serum immunoglobulins, particularly IgM, IgG2b, and IgG3. This upregulation could be the result of increased B cell metabolism, a change in antigen exposure, and/or related to increased innate-like B cell subsets. Second, both Lean and DIO SG had increased splenic and intestinal Ly6G^+^CD11B^+^ neutrophils. Finally, there was a small but statistically significant increase in splenic and intestinal Arg1^+^ M2 macrophages. Together, these represent one of the first multi-organ characterization of the immune effects of SG in weight-loss dependent and weight loss independent models.

There is an abundance of literature highlighting the effects of obesity on global and organ specific immune cell changes and their relation to T2D. Studies in DIO mice have established roles for chronic inflammation within the visceral adipose tissue, hepatic, and intestine in the pathogenesis of obesity induced T2D. Multiple cell-specific changes associated with DIO have been documented including enlargement of the macrophage and DC pools, CD8^+^ and Th1 T cell recruitment, and decreased regulatory T cell populations.(6,11,35–37) We demonstrate that surgery alters immune subsets independent of weight loss and raise the possibility that these changes contribute to the improvements in insulin sensitivity seen post-surgery.

There is strong evidence from the DIO literature that each of the key immune subsets we find altered by SG play a role in glucose metabolism. A specific role for B lymphocytes in the pathogenesis of T2DM in murine DIO models has been demonstrated. Winer et al. show that B-2 cells, a subtype that includes mature follicular B cells, in DIO animals undergo class switching to IgG, produce less anti-inflammatory IL-10, and produce more pro-inflammatory IL-8. B cell deficient DIO mice exhibit better glycemic control than wildtype animals. Adoptive transfer of either B-2 cells or IgG from DIO animals to B^null^ animals worsens metabolic disease.(8,38) Further, B cells have been shown to support pro-inflammatory T cell function in T2D through contact dependent mechanisms.(12) Our findings of reduced splenic CD19^+^CD11b^neg^ B cells post-SG would be predicted to contribute to improvement in insulin resistance.

On the other hand, several populations of innate-like B cells – including B1a, B-reg, and innate B cells – improve insulin sensitivity via production of IL-10 and/or IgM.(13,14,17,39) These findings have been confirmed in humans, where a deficiency in IL-10 production by peripheral B cells is a conserved feature of diabetes.(40) In the spleen, we find that SG increases CD11b^+^ B cells, which may represent an innate-like B cell phenotype.(16) Innate B cells are predominantly located in the peritoneal cavity, adipose tissue, and spleen.(13–16) Unfortunately, we did not include adipose depots in our organs for evaluation owing to the relatively low yield in lean sham and SG mice. Frikke-Schmidt et al., evaluated adipose immune subsets post-SG; however, they did not include B cell markers in their analysis.(22) In humans, Roux-en-Y gastric bypass (RYGB) leads to increased circulating IL-10 producing B cells and SG has been associated with elevated levels of IgG, IgA, IgM up to 12 months surgery.(21,41) The effect of SG on innate, B1-like population frequency and function and its consequences for glucose metabolism merits future evaluation.

We also observed weight-independent effects on myeloid cells, including neutrophils and macrophages. We find a conserved significant but small increase in Arg1 expression across multiple macrophage populations in Lean and DIO SG mice, suggestive of increased M2 polarization. There is also a nearly 5.5-fold increase in splenic neutrophils. Imbalances in macrophage polarization have been implicated in a number of inflammatory diseases, including obesity and T2D.(42) M2 polarization, particularly in adipose tissue, improves insulin sensitivity. Two prior studies showed increased neutrophil recruitment to subcutaneous adipose tissue in human subjects with obesity, with and without T2D, after SG or RYGB as compared to their presurgical specimen.(43,44) Neutrophils, while classically are innate responders to blood borne pathogens, also take residence in the marginal zone (MZ) of the spleen in response to gut commensal organisms via non-inflammatory pathways. These B-helper neutrophils (N_BH_) orchestrate a complex interaction with MZ, innate-like B cells to promote class switching, somatic hypermutation, and the production of IgG, IgM, and IgA.(45,46) Similarly, crosstalk between M1/M2 macrophages and both B2 and innate B1-like B cells influences insulin resistance.(12) More detailed studies will be needed to understand whether SG induced changes in neutrophils, macrophages, and B cells are interconnected and their consequence on insulin resistance.

The mechanisms by which SG alters cellular immune populations needs of further investigation. Nutrient availability and immunometabolism may affect immune function. B cells are sensitive to their environment and changes in nutrient availability influences their differentiation.(47) Diet can change the gut microbial community and intestinal homeostasis/barrier function leading to translocation of novel antigens and adipose inflammation.(35,48) Thus, new antigens may be exposed or created after SG triggering immune modulation. Immune cells also express bile acid and incretin hormone receptors, such as GLP-1. We have previously shown that DIO SG animals have increased levels of a bile acid metabolite, cholic acid 7-sulfate, which is a potent agonist of Takeda g-protein bile acid receptor (TGR5).(31) Thus, SG-alterations in local and systemic BA and incretin hormones may have consequences on immunity.(49)

Our study has several limitations. Our screening design is potentially underpowered to detect differences in rare immune sub-populations. We focused on cell surface markers to phenotype cellular immune populations. As such, we may have missed changes in immune cell function that occurred without changes in frequency. As mentioned previously, due to cell numbers, we focused on splenic, hepatic, and intestinal immune populations and did not include adipose depots or peritoneal cavity organs in this study, which is an area that needs future study. Future studies will be needed to understand the metabolic and immunologic consequences of SG induced changes in immunity and their driving mechanisms. It will also be essential to correlate these findings in blood and tissue from human patients to understand their clinical relevance.

## Conclusions

SG leads to a conserved metabolic and immunologic phenotype independent of weight loss. CyTOF profiling identified a shift in the splenic B cell compartment from follicular to innate-like B cells in DIO SG mice with a trend toward a similar shift in Lean SG animals. These changes were associated with an increase in total IgM, total IgG, and IgG2b in serum. Further, SG lead to an increase in splenic neutrophils and M2 macrophage polarization. These weight-loss independent, surgery-specific immune changes have been associated with improvement in glucose metabolism and have the potential to be contributors to post SG physiology.

## Acknowledgements

Experiments were designed and interpreted by DAH, RS, TB, AT, EGS. Surgical procedures, animal care, and functional experiments were completed by DAH, TB, and RS. E.G.S. is the guarantor of this work and, as such, had full access to all the data in the study and takes responsibility for the integrity of the data and the accuracy of the data analysis. We would like to thank the Harvard Medical Area CyTOF Antibody and Resource Core and Dana Farber Cancer Institute, Longwood Medical Area CyTOF Core facility for their invaluable help in developing this project.

This work was conducted with the support of funding sources from the senior and first author. First, was an appointed KL2 award from Harvard Catalyst. The Harvard Clinical and Translational Science Center (National Center for Advancing Translational Sciences, National Institutes of Health Award KL2 TR002542). The content is solely the responsibility of the authors and does not necessarily represent the official views of Harvard Catalyst, Harvard University and its affiliated academic healthcare centers, or the National Institutes of Health. Second, funding was obtained through a Pilot Grant from the Boston Area Diabetes and Endocrinology Research Center (BADERC – NIH P30 DK057521). A third funding source was the New England Surgical Society Scholars Foundation Research Grant. DAH received a two-year, competitive research scholarship from the American College of Surgeons and was funded, in part, through an NIH T32 (T32DK007754) during the time this work was completed.

## Abbreviations

SG: (Sleeve Gastrectomy)
(T2D): Type 2 Diabetes
(TNF-α): tumor necrosis factor alpha
(VAT): visceral adipose tissue
POD: (post-operative Day)
GLP-1: (glucagon like peptide 1)
OGTT: (oral glucose tolerance test)
ITT: (insulin tolerance test)
DIO: (diet induced obesity)
CyTOF: (cytometry with time of flight analysis)
ARG1: (arginase 1)
TGR5: (Takeda g-protein bile acid receptor)
RYGB: (Roux-en-Y gastric bypass)

