## Supplementary figures and images for "Weight and organ specific immune cell profiling of Sleeve Gastrectomy"

### Supplemental Figures

#### Supplemental Figures

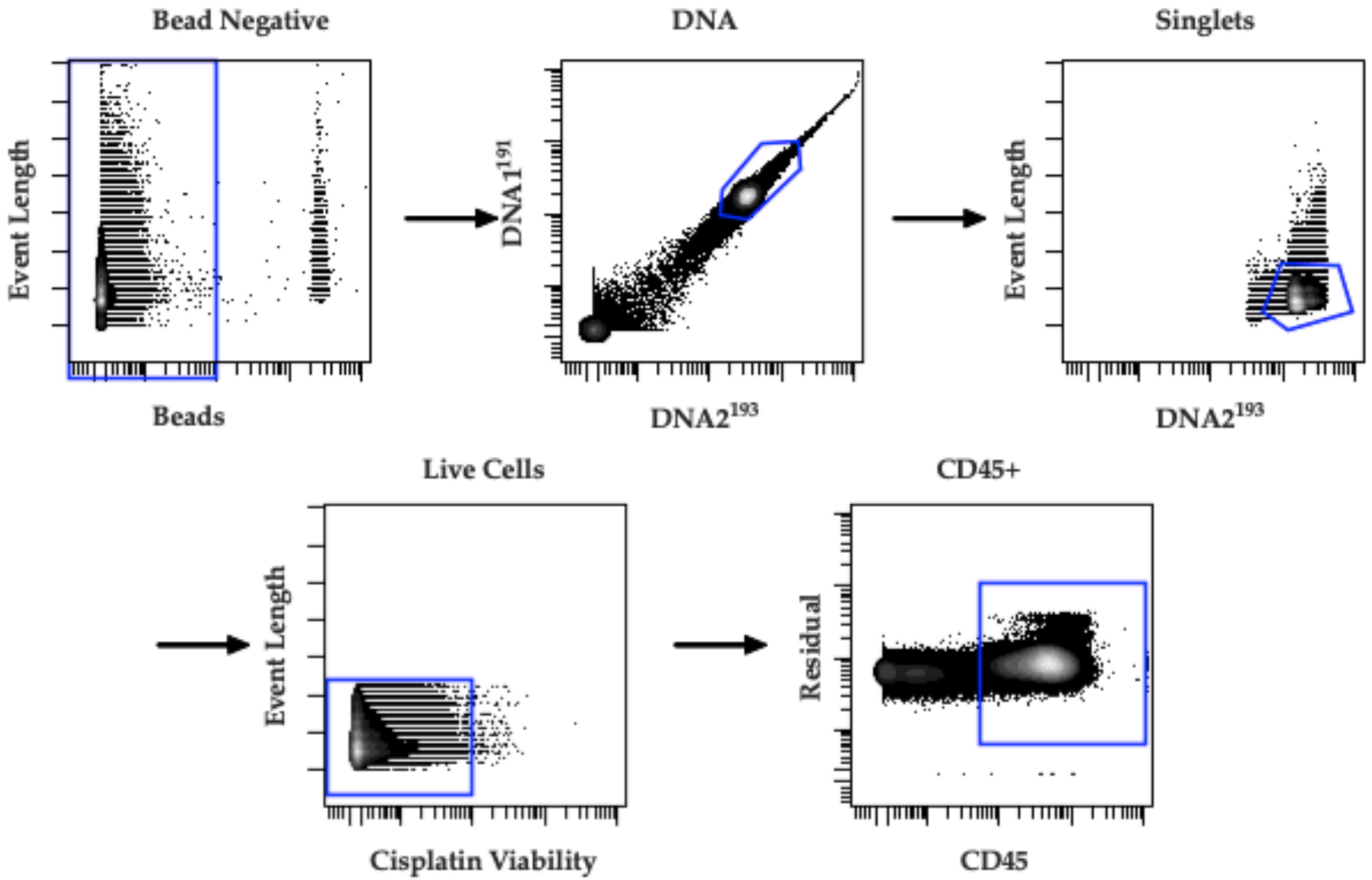


**Supp figure 1**

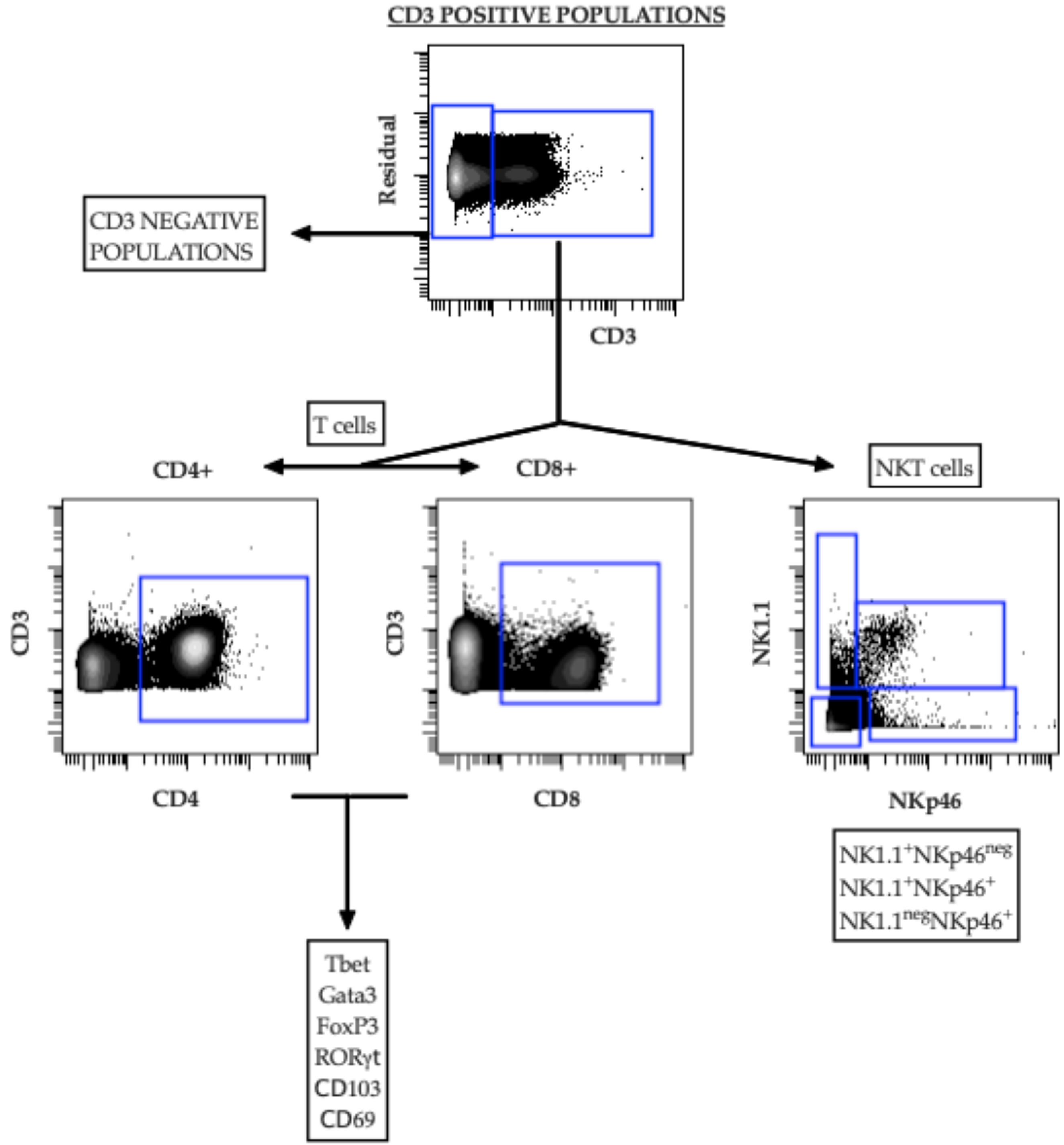


**Supp figure 2**

Supp
Figure 3

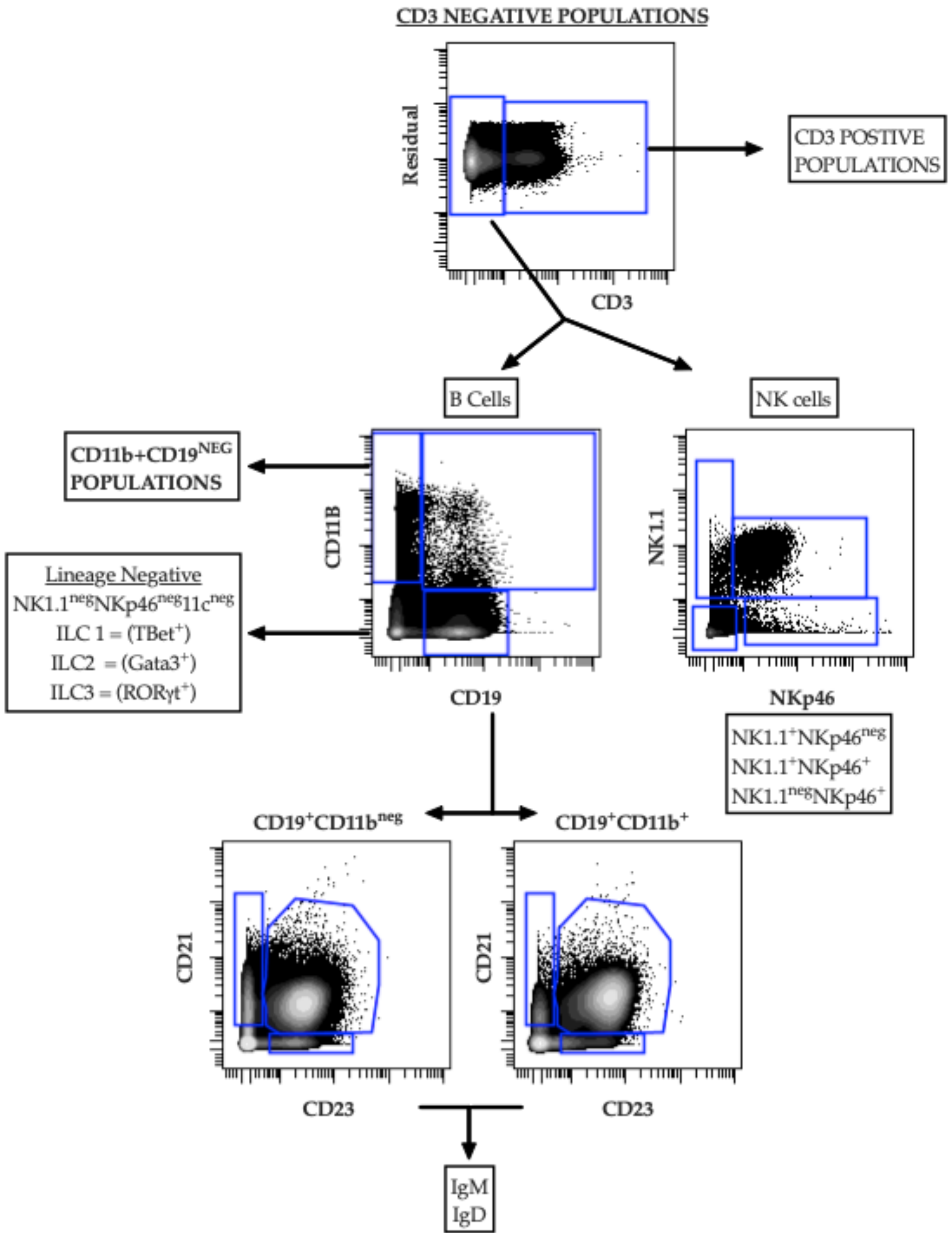

**Supp Figure 4**

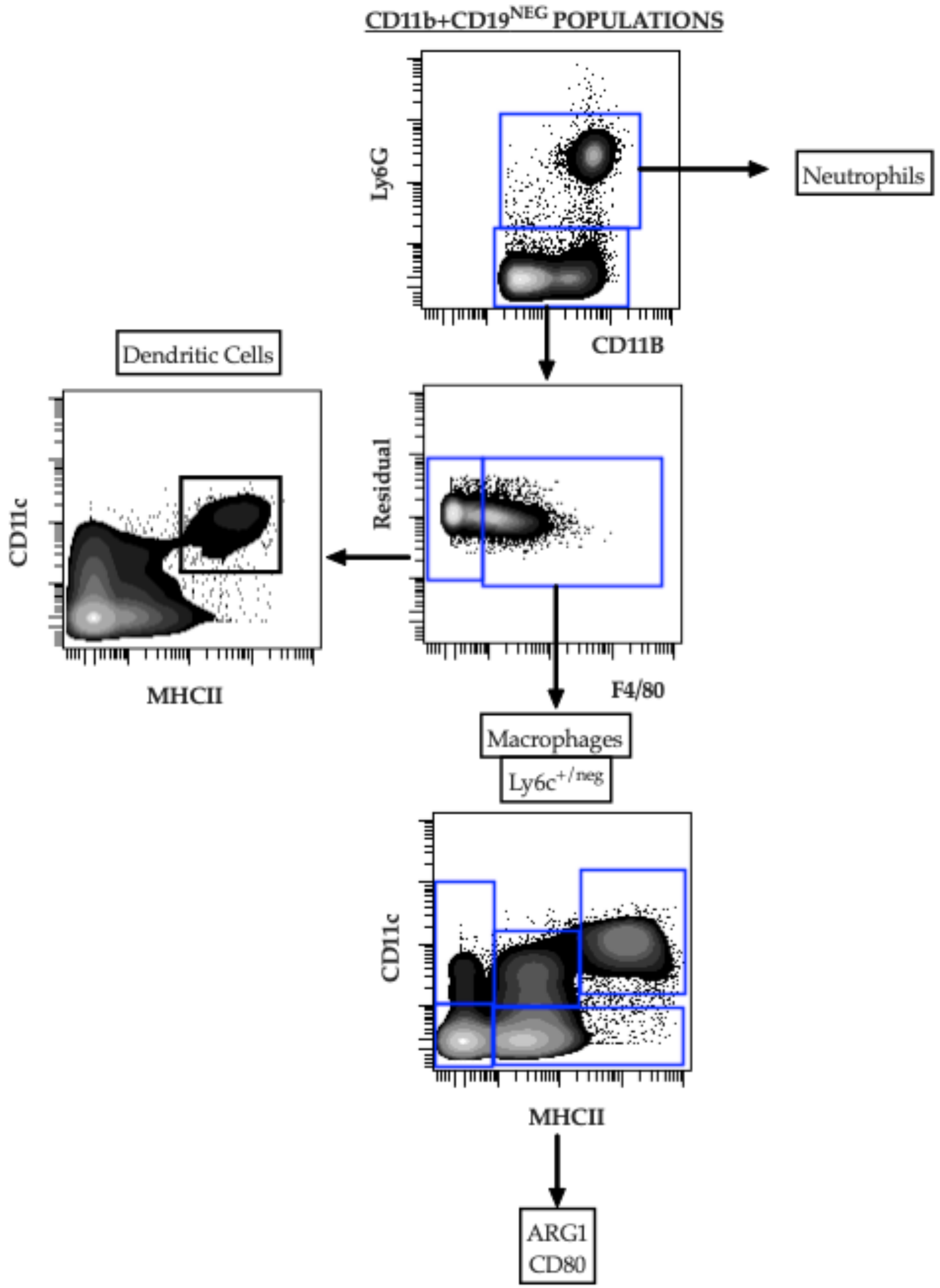
