## Supplemental Tables for "Weight and organ specific immune cell profiling of Sleeve Gastrectomy"

| **Supplemental Table 1. CyTOF Panel** | | |
| --- | --- | --- |
| **Antibody Target** | **Metal Conjugate** | **Clone** |
| CD45 | 141Pr | 30-F11 |
| CD21/35 | 143Nd | 2e9 |
| CD5 | 144Nd | 53-7.3 |
| CD4 | 145Nd | RM4-5 |
| CD11c | 146Nd | N418 |
| Ly6G | 148nd | 1A8 |
| CD19 | 149Sm | 6D5 |
| LY6C | 151Eu | HK1.4 |
| CD3 | 152Sm | 145-2C11 |
| CD335/NKp46 | 153Eu | 29A1.4 |
| *CD80 | 154Sm | 16-10A1 |
| *IgM | 156gd | rmm1 |
| T-bet | 158gd | 4B10 |
| CD23 | 159tb | B3B4 |
| CD11b | 160Gd | M1/70 |
| Arginase 1 | 161DY | polyclonal |
| FoxP3 | 162Dy | FJK-163 |
| NK1.1 | 163DY | PK136 |
| CD8a | 164DY | 53-6.7 |
| CD103 | 166Er | 2E7 |
| CD25/IL-2Rα | 174Yb | 3C7 |
| CD64 | 168Er | x54-5/7.1 |
| CCR6 | 169Tm | 29-2L17 |
| ROR-γt | 170Er | AFKJS-9 |
| GATA3 | 172Yb | TWAJ |
| CD69 | 173YB | H1.2F3 |
| F4/80 | 174YB | BM8 |
| IgD | 175Lu | 11-26c.2a |
| MHCII (IA/IE) | 209Bi | M5/114.15.2 |

*Antibodies not included in the first panel iteration, which was used in the first Lean and DIO experiments

| **Supplemental Table 2: T cell subsets as % of CD45+ Leukocytes – Lean Sham versus SG** | | | | | | | | | | | | | | | | | | | |  |  |  |
| --- | --- | --- | --- | --- | --- | --- | --- | --- | --- | --- | --- | --- | --- | --- | --- | --- | --- | --- | --- | --- | --- | --- |
|  | **Jejunum** | | | | **Ileum** | | | | | **Cecum** | | | | | **Spleen** | | | | | **Liver** | | |
|  | Sham | SG | | p | Sham | SG | | | P | Sham | SG | | | p | Sham | | SG | | p | Sham | SG | P |
| CD3+ | 30.1±9.6 | 33.3±19.1 | 0.644 | | 45.7±5.3 | | 28.9±12.9 | 0.076 | | 59.6±19.3 | | 55.7±11.8 | 0.704 | | 25.8±2.9 | 22.7±7.4 | | 0.235 | | 19.3±7.3 | 15.9±3.1 | 0.415 |
| CD3+103+ | 12.3±7.2 | 15.3±15.9 | 0.597 | | 3.8±1.9 | | 3.8±2.6 | 0.976 | | 8.5±8.1 | | 9.5±2.4 | 0.809 | | 2.9±1.1 | 2.2±1.5 | | 0.248 | | 0.90±0.68 | 0.57±0.23 | 0.398 |
| CD3+CD69+ | 12.0±5.1 | 14.2±10.3 | 0.560 | | 16.1±11.6 | | 10.5±6.8 | 0.221 | | 35.9±16.1 | | 33.4±11.3 | 0.789 | | 2.4±0.91 | 2.3±1.0 | | 0.706 | | 7.1±6.1 | 4.7±4.2 | 0.523 |
| CD4+ | 3.2±1.3 | 3.4±2.0 | 0.864 | | 11.7±6.4 | | 10.8±4.5 | 0.730 | | 6.6±2.6 | | 9.1±1.4 | 0.098 | | 14.5±2.3 | 13.2±4.9 | | 0.461 | | 11.2±5.2 | 8.0±0.9 | 0.272 |
| CD4+CD103+ | 0.72±0.47 | 0.64±0.23 | 0.639 | | 0.49±0.41 | | 0.88±0.65 | 0.140 | | 0.27±0.18 | | 0.43±0.10 | 0.136 | | 0.28±0.12 | 0.27±0.13 | | 0.817 | | 0.21±0.11 | 0.15±0.09 | 0.435 |
| CD4+FOXP3+ | 0.43±0.36 | 0.41±0.38 | 0.913 | | 2.0±2.0 | | 1.9±2.0 | 0.933 | | 0.11±0.04 | | 0.14±0.06 | 0.307 | | 0.31±0.17 | 0.28±0.18 | | 0.692 | | 1.2±2.1 | 0.23±0.26 | 0.400 |
| CD4+GATA3+ | 0.52±0.48 | 0.71±0.85 | 0.555 | | 3.6±5.2 | | 2.5±2.9 | 0.581 | | 0.11±0.14 | | 0.22±0.20 | 0.338 | | 0.91±0.6 | 0.74±0.9 | | 0.629 | | 3.5±5.1 | 1.6±1.6 | 0.487 |
| CD4+RORγt+ | 0.34±0.25 | 0.42±0.45 | 0.602 | | 2.1±2.7 | | 1.9±2.4 | 0.901 | | 0.05±0.04 | | 0.12±0.08 | 0.151 | | 0.05±0.03 | 0.03±0.02 | | 0.071 | | 0.88±1.61 | 0.11±0.13 | 0.380 |
| CD4+Tbet+ | 0.14±0.11 | 0.31±0.37 | 0.168 | | 0.54±0.68 | | 0.84±1.9 | 0.646 | | 0.10±0.06 | | 0.16±0.13 | 0.344 | | 0.19±0.19 | 0.12±0.14 | | 0.367 | | 0.03±0.03 | 0.01±0.01 | 0.286 |
| CD8+ | 14.7±9.4 | 16.0±18.6 | 0.854 | | 5.5±2.5 | | 5.8±3.0 | 0.814 | | 9.9±6.2 | | 11.8±1.7 | 0.518 | | 9.9±1.2 | 8.3±2.9 | | 0.135 | | 4.5±1.2 | 4.6±0.8 | 0.892 |
| CD8+CD103+ | 8.7±6.2 | 9.9±13.8 | 0.808 | | 2.0±1.3 | | 2.2±2.0 | 0.788 | | 4.2±4.2 | | 4.6±1.8 | 0.822 | | 2.5±1.0 | 1.8±1.4 | | 0.218 | | 0.67±0.55 | 0.35±0.11 | 0.301 |
| CD8+CD103+CD69+ | 4.9±3.6 | 5.4±7.1 | 0.846 | | 0.44±0.51 | | 0.89±1.1 | 0.249 | | 3.1±3.5 | | 3.4±1.5 | 0.876 | | 0.09±0.05 | 0.07±0.06 | | 0.580 | | 0.03±0.02 | 0.02±0.02 | 0.301 |
| CD8+FoxP3+ | 0.30±0.22 | 0.33±0.27 | 0.754 | | 0.72±0.92 | | 0.91±1.2 | 0.702 | | 0.09±0.06 | | 0.13±0.09 | 0.488 | | 0.05±0.02 | 0.04±0.02 | | 0.334 | | 0.49±0.76 | 0.08±0.10 | 0.438 |
| CD8+GATA3+ | 0.51±0.30 | 0.76±0.59 | 0.251 | | 0.94±1.3 | | 1.1±1.3 | 0.754 | | 0.14±0.09 | | 0.24±0.15 | 0.256 | | 0.36±0.21 | 0.33±0.45 | | 0.844 | | 0.79±1.2 | 0.54±0.59 | 0.710 |
| CD8+Tbet+ | 1.4±2.2 | 3.2±7.0 | 0.452 | | 0.83±1.2 | | 0.93±1.4 | 0.870 | | 1.1±1.0 | | 1.1±0.45 | 0.992 | | 0.65±0.71 | 0.51±0.57 | | 0.638 | | 0.01±0.02 | 0.00±0.01 | 0.288 |

Average with SD. Students T-test. P< 0.05 highlighted. Jejunal, Ileal, and splenic specimen with n 10 (sham), 9 (SG). Data from these specimens represent combined results of two replicate experiments. Liver specimen with n 5 (sham), 4 (SG). Cecal samples with n 5 (sham), 5 (SG).

| **Supplemental Table 3: NKT, NK, IL cell subsets as % of CD45+ Leukocytes – Lean Sham versus SG** | | | | | | | | | | | | | | | | | | | | |  |  |  |
| --- | --- | --- | --- | --- | --- | --- | --- | --- | --- | --- | --- | --- | --- | --- | --- | --- | --- | --- | --- | --- | --- | --- | --- |
|  | **Jejunum** | | | | | **Ileum** | | | | | **Cecum** | | | | | **Spleen** | | | | | **Liver** | | |
|  | Sham | SG | | | p | Sham | Sham | | | P | Sham | SG | | | p | Sham | | SG | | p | Sham | SG | P |
| CD3+NK1.1+ | 0.11±0.11 | | 0.11±0.13 | 0.974 | | 0.38±0.61 | | 0.08±0.09 | 0.158 | | 2.1±1.6 | | 1.4±0.6 | 0.432 | | 0.22±0.12 | 0.20±0.12 | | 0.729 | | 1.1±1.4 | 0.75±0.90 | 0.647 |
| CD3+NKp46+ | 0.90±0.58 | | 0.82±0.80 | 0.819 | | 4.1±2.5 | | 2.3±2.1 | 0.291 | | 1.3±0.5 | | 1.1±0.5 | 0.577 | | 0.29±0.13 | 0.20±0.10 | | 0.122 | | 0.28±0.22 | 0.29±0.28 | 0.964 |
| CD3+NK1.1+NKp46+ | 0.15±0.10 | | 0.18±0.18 | 0.702 | | 0.37±0.47 | | 0.61±1.1 | 0.532 | | 0.29±0.26 | | 0.24±0.08 | 0.712 | | 0.15±0.08 | 0.14±0.07 | | 0.886 | | 0.11±0.10 | 0.13±0.20 | 0.810 |
| CD3^neg^NK1.1+ | 0.22±0.24 | | 0.18±0.12 | 0.693 | | 0.1±0.09 | | 0.13±0.11 | 0.595 | | 1.3±1.4 | | 1.4±0.2 | 0.961 | | **0.32±0.14** | **0.65±0.36** | | **0.016** | | 1.5±0.8 | 1.1±0.4 | 0.372 |
| CD3^neg^NKp46+ | 1.4±0.6 | | 1.3±1.2 | 0.850 | | 2.3±1.9 | | 2.8±2.0 | 0.609 | | 0.71±0.51 | | 0.73±0.25 | 0.919 | | 0.85±0.21 | 1.0±0.4 | | 0.806 | | 1.2±0.5 | 2.1±1.7 | 0.281 |
| CD3^neg^NK1.1+NKp46+ | 0.06±0.04 | | 0.09±0.06 | 0.210 | | 0.06±0.06 | | 0.06±0.05 | 0.977 | | **0.41±0.26** | | **1.1±0.6** | **0.039** | | 2.0±0.6 | 2.0±0.8 | | 0.806 | | 1.7±1.0 | 1.9±1.5 | 0.826 |
| ILC1* (TBet+) | 0.31±0.46 | | 0.38±0.49 | 0.758 | | 0.09±0.10 | | 0.27±0.29 | 0.089 | | 0.56±0.35 | | 0.65±0.13 | 0.594 | | 0.20±0.13 | 0.21±0.22 | | 0.895 | | 0.06±0.04 | 0.02±0.02 | 0.143 |
| ILC2* (Gata3+) | 0.51±0.45 | | 0.66±0.49 | 0.496 | | 0.61±0.51 | | 1.2±1.3 | 0.185 | | 0.19±0.10 | | 0.34±0.35 | 0.383 | | 0.83±0.75 | 3.3±8.4 | | 0.372 | | 3.4±3.8 | 2.2±1.5 | 0.560 |
| ILC3* (RORγt+) | 0.07±0.03 | | 0.09±0.07 | 0.502 | | 0.10±0.11 | | 0.13±0.13 | 0.604 | | 0.03±0.01 | | 0.02±0.02 | 0.324 | | 0.04±0.02 | 0.03±0.04 | | 0.612 | | 0.42±0.62 | 0.08±0.05 | 0.318 |

* Innate Lymphoid Cell. Derivative of a lineage negative population (CD45+CD3^neg^CD19^neg^CD11b^neg^CD11c^neg^NK1.1^neg^NKp46^neg^). Average with SD. Students T-test. P< 0.05 highlighted. Jejunal, Ileal, and splenic specimen with n 10 (sham), 9 (SG). Data from these specimens represent combined results of two replicate experiments. Liver specimen with n 5 (sham), 4 (SG). Cecal samples with n 5 (sham), 5 (SG).

| **Supplemental Table 4: CD19+11B**^neg^ **B Cell subsets as % of CD45+ Leukocytes – Lean Sham versus SG** | | | | | | | | | | | | | | | | | | |  |  |  |
| --- | --- | --- | --- | --- | --- | --- | --- | --- | --- | --- | --- | --- | --- | --- | --- | --- | --- | --- | --- | --- | --- |
|  | **Jejunum** | | | **Ileum** | | | | | **Cecum** | | | | | **Spleen** | | | | | **Liver** | | |
|  | Sham | SG | p | Sham | SG | | | p | Sham | SG | | | p | Sham | | SG | | p | Sham | SG | P |
| Total | 4.5±3.8 | 3.8±8.0 | 0.815 | 19.7±13.7 | | 24.2±16.1 | 0.521 | | 23.5±16.7 | | 22.1±10.4 | 0.877 | | **62.0±4.7** | **47.1±19.5** | | **0.031** | | 28.8±14.2 | 24.4±14.9 | 0.669 |
| #Total CD80+ | 0.12±0.10 | 0.09±0.09 | 0.725 | 0.46±0.37 | | 0.95±0.56 | 0.143 | | 0.27±0.24 | | 0.31±0.12 | 0.753 | | 1.3±0.4 | 1.4±0.6 | | 0.600 | |  | NA |  |
| #Total IgM^neg^IgD+ | 1.1±1.0 | 0.21±0.018 | 0.081 | 3.9±3.7 | | 5.3±3.3 | 0.535 | | 6.5±4.9 | | 5.3±2.7 | 0.643 | | 6.7±2.0 | 8.1±1.8 | | 0.268 | |  | NA |  |
| #Total IgM+IgD+ | 4.0±3.2 | 0.79±0.81 | 0.061 | 12.6±11.8 | | 20.1±16.3 | 0.428 | | 14.7±10.7 | | 13.9±7.8 | 0.897 | | **54.0±1.84** | **40.5±10.2** | | **0.020** | |  | NA |  |
| #Total IgM+IgD^neg^ | 0.25±0.28 | 0.12±0.07 | 0.327 | 0.91±0.98 | | 1.6±0.6 | 0.218 | | 0.47±0.30 | | 0.82±0.28 | 0.093 | | 3.5±0.9 | 4.0±1.2 | | 0.505 | |  | NA |  |
| Total CD21+CD23^neg^ | 0.86±1.2 | 0.25±0.30 | 0.175 | 3.2±4.7 | | 4.8±5.5 | 0.499 | | 5.8±4.0 | | 5.3±2.3 | 0.841 | | 6.3±3.1 | 6.9±3.9 | | 0.713 | | 0.58±0.34 | 0.87±0.97 | 0.547 |
| Total CD21+CD23+ | 0.27±0.30 | 0.20±0.41 | 0.711 | 0.75±0.93 | | 1.3±1.5 | 0.306 | | 10.9±8.3 | | 9.0±5.0 | 0.672 | | **41.8±6.4** | **28.4±13.0** | | **0.010** | | 10.2±8.3 | 8.7±6.8 | 0.789 |
| Total CD21^neg^CD23+ | 1.4±1.0 | 1.8±4.6 | 0.787 | 6.4±5.1 | | 7.6±7.3 | 0.663 | | 1.6±1.3 | | 1.9±1.2 | 0.674 | | 1.6±0.6 | 2.3±1.5 | | 0.153 | | 0.36±0.17 | 0.88±0.83 | 0.212 |
| #CD21+CD23+IgM^neg^IgD+ | 0.26±0.24 | 0.06±0.08 | 0.116 | 0.96±0.81 | | 1.7±1.5 | 0.349 | | 2.9±2.3 | | 2.1±1.2 | 0.511 | | 5.1±1.4 | 5.6±1.7 | | 0.598 | |  | NA |  |
| #CD21+CD23+IgM+IgD- | 0.03±0.03 | 0.01±0.02 | 0.539 | 0.08±0.11 | | 0.13±0.09 | 0.511 | | 0.07±0.06 | | 0.08±0.05 | 0.756 | | 0.34±0.10 | 0.49±0.17 | | 0.121 | |  | NA |  |
| #CD21+CD23+IgM+IgD+ | 1.3±1.0 | 0.26±0.33 | 0.058 | 4.1±4.0 | | 7.6±8.0 | 0.408 | | 7.7±5.9 | | 6.6±3.9 | 0.753 | | **39.4±3.8** | **27.9±8.3** | | **0.022** | |  | NA |  |

All B cell subsets from CD19+CD11b^neg^ parent. Average with SD. Students T-test. P< 0.05 highlighted. Jejunal, Ileal, and splenic specimen with n 10 (sham), 9 (SG) except as denoted by the symbol, “#”. Data from these specimens represent combined results of two replicate experiments. Liver specimen with n 5 (sham), 4 (SG) and for this experiment, IgD and CD80 were not in the staining panel (NA). Cecal samples with n 5 (sham), 5 (SG). #: data is strictly from the second panel iteration that included IgD and CD80 (n 5,5).

| **Supplemental Table 5: CD19+11B+ B Cell subsets as % of CD45+ Leukocytes – Lean versus SG** | | | | | | | | | | | | | | | | | | |  |  | |  |
| --- | --- | --- | --- | --- | --- | --- | --- | --- | --- | --- | --- | --- | --- | --- | --- | --- | --- | --- | --- | --- | --- | --- |
|  | **Jejunum** | | | **Ileum** | | | | | **Cecum** | | | | | **Spleen** | | | | | **Liver** | | | |
|  | Sham | SG | p | Sham | | SG | | p | Sham | | SG | | p | Sham | | SG | p | | Sham | SG | | P |
| Total | 0.14±0.05 | 0.10±0.06 | 0.298 | 0.04±0.04 | 0.17±0.21 | | 0.226 | | 0.05±0.06 | 0.05±0.04 | | 0.872 | | 0.84±0.21 | 2.1±2.4 | | | 0.274 | 0.28±0.33 | 0.72±1.1 | 0.405 | |
| #Total CD80+ | 0.04±0.03 | 0.07±0.05 | 0.285 | 0.03±0.05 | 0.06±0.07 | | 0.390 | | 0.02±0.02 | 0.02±0.02 | | 0.819 | | 0.22±0.06 | 0.39±0.30 | | | 0.262 |  | NA | |  |
| #Total IgM^neg^IgD+ | **0.02±0.01** | **0.00±0.00** | **0.010** | 0.01±0.03 | 0.00±0.00 | | 0.347 | | 0.01±0.02 | 0.02±0.03 | | 0.640 | | 0.08±0.05 | 0.26±0.31 | | | 0.248 |  | NA | |  |
| #Total IgM+IgD+ | 0.05±0.04 | 0.03±0.02 | 0.353 | 0.01±0.02 | 0.09±0.15 | | 0.267 | | 0.03±0.03 | 0.03±0.02 | | 0.984 | | 0.54±0.13 | 1.4±1.8 | | | 0.298 |  | NA | |  |
| #Total IgM+IgD^neg^ | 0.02±0.03 | 0.03±0.03 | 0.783 | 0.01±0.01 | 0.05±0.07 | | 0.268 | | 0.00±0.01 | 0.01±0.01 | | 0.728 | | 0.10±0.03 | 0.18±0.13 | | | 0.174 |  | NA | |  |
| Total CD21+CD23^neg^ | 0.02±0.02 | 0.05±0.05 | 0.330 | 0.01±0.01 | 0.06±0.07 | | 0.159 | | 0.02±0.03 | 0.01±0.01 | | 0.778 | | 0.18±0.06 | 0.38±0.39 | | | 0.287 | 0.001±0.01 | 0.01±0.01 | | 0.448 |
| Total CD21^neg^CD23+ | 0.01±0.01 | 0.00±0.01 | 0.611 | 0.00±0.00 | 0.00±0.00 | | 0.347 | | 0.00±0.00 | 0.00±0.01 | | 0.529 | | 0.02±0.02 | 0.12±0.19 | | | 0.258 | 0.09±0.16 | 0.20±0.34 | | 0.539 |
| Total CD21+CD23+ | 0.03±0.02 | 0.01±0.01 | 0.072 | 0.02±0.03 | 0.04±0.08 | | 0.500 | | 0.02±0.02 | 0.02±0.02 | | 0.965 | | 0.33±0.15 | 1.1±1.3 | | | 0.240 | 0.004±0.01 | 0.02±0.04 | | 0.368 |

All B cell subsets from CD19+CD11b+ parent. Average with SD. Students T-test. P< 0.05 highlighted. Jejunal, Ileal, and splenic specimen with n 10 (sham), 9 (SG) except as denoted by the symbol, “#”. Data from these specimens represent combined results of two replicate experiments. Liver specimen with n 5 (sham), 4 (SG) and for this experiment, IgD and CD80 were not in the staining panel (NA). Cecal samples with n 5 (sham), 5 (SG). #: data is strictly from the second panel iteration that included IgD and CD80 (n 5,5).

| **Supplemental Table 6: Myeloid Cell subsets as % of CD45+ Leukocytes – Lean versus SG** | | | | | | | | | | | | | |  |  |  |
| --- | --- | --- | --- | --- | --- | --- | --- | --- | --- | --- | --- | --- | --- | --- | --- | --- |
|  | **Jejunum** | | | **Ileum** | | | **Cecum** | | | **Spleen** | | | | **Liver** | | |
|  | Sham | SG | p | Sham | SG | p | Sham | SG | p | Sham | SG | | p | Sham | SG | P |
| ***PMN***: Ly6G+ CD11b+ | 0.32±0.28 | 0.37±0.30 | 0.696 | **0.09±0.12** | **0.27±0.17** | **0.014** | 0.14±0.06 | 0.38±0.45 | 0.273 | **1.3±0.7** | **7.1±6.3** | **0.009** | | 6.0±2.0 | 13.4±12.6 | 0.231 |
| ***DC:***  F480 ^neg^11C+MHCII+ | 0.14±0.09 | 0.19±0.15 | 0.330 | 0.06±0.04 | 0.09±0.06 | 0.322 | 0.08±0.07 | 0.08±0.02 | 0.945 | **0.23±0.09** | **0.72±0.61** | **0.022** | | 0.42±0.18 | 0.50±0.12 | 0.523 |
| ***MACS***  ***(Ly6GG*** ^neg^ ***F480+)*** |  |  |  |  |  |  |  |  |  |  |  |  | |  |  |  |
| LY6C+ | 1.13±0.82 | 0.90±0.62 | 0.513 | 0.10±0.10 | 0.30±0.37 | 0.123 | 0.18±0.14 | 0.22±0.12 | 0.685 | 0.50±0.31 | 1.6±2.1 | 0.142 | | 1.2±0.7 | 2.5±3.4 | 0.423 |
| LYGC ^neg^ | 17.7±8.0 | 14.9±10.2 | 0.517 | **0.45±0.28** | **1.2±0.8** | **0.011** | 0.38±0.36 | 0.28±0.15 | 0.585 | 0.72±0.30 | 2.5±3.6 | 0.133 | | 1.6±0.8 | 3.3±3.6 | 0.316 |
| CD11C+MHCII+ | 0.29±0.22 | 0.51±0.56 | 0.273 | 0.14±0.16 | 0.25±0.45 | 0.468 | 0.16±0.13 | 0.10±0.03 | 0.326 | 0.41±0.21 | 0.66±0.55 | 0.207 | | 0.49±0.26 | 0.73±0.73 | 0.515 |
| CD11C+MHClo | 2.5±1.1 | 1.6±0.83 | 0.077 | **0.06±0.06** | **0.18±012** | **0.016** | 0.01±0.01 | 0.00±0.01 | 0.672 | 0.17±0.15 | 0.56±0.92 | 0.210 | | 0.38±0.52 | 1.0±1.6 | 0.421 |
| CD11C ^neg^MHCII+ | 0.71±0.45 | 0.72±0.70 | 0.958 | 0.15±0.14 | 0.28±0.25 | 0.178 | 0.18±0.15 | 0.17±0.12 | 0.926 | 0.26±0.25 | 1.6±3.8 | 0.276 | | 0.52±0.37 | 1.3±1.9 | 0.368 |
| CD11C ^neg^MHCII^neg^ | 2.7±1.8 | 3.0±2.6 | 0.823 | **0.10±0.09** | **0.42±0.3** | **0.006** | 0.19±0.21 | 0.18±0.08 | 0.888 | 0.21±0.13 | 0.73±0.81 | 0.061 | | 0.66±0.08 | 1.2±1.1 | 0.289 |
| CD11C+MHCII^neg^ | 12.8±8.1 | 10.2±7.7 | 0.475 | 0.09±0.09 | 0.37±0.42 | 0.058 | 0.02±0.01 | 0.05±0.04 | 0.225 | **0.13±0.10** | **0.38±0.31** | **0.029** | | 0.53±0.29 | 1.1±1.2 | 0.328 |

All Macrophage populations are F480+. Average with SD. Students T-test. P< 0.05 highlighted. Jejunal, Ileal, and splenic specimen with n 10 (sham), 9 (SG). Data from these specimens represent combined results of two replicate experiments. Liver specimen with n 5 (sham), 4 (SG). Cecal samples with n 5 (sham), 5 (SG). Ly6C expression was measured from total macrophage parent

| **Table 7: CD80+ M1 Macrophage polarization as percent of Parent – Lean Sham versus SG** | | | | | | | | | | | | | |  |  |  |
| --- | --- | --- | --- | --- | --- | --- | --- | --- | --- | --- | --- | --- | --- | --- | --- | --- |
|  | **Jejunum** | | | **Ileum** | | | **Cecum** | | | **Spleen** | | | | **Liver** | | |
|  | Sham | SG | p | Sham | SG | p | Sham | SG | p | Sham | SG | | p | Sham | SG | P |
| MACS: Ly6G^neg^F480+ | 65.6±8.8 | 66.8±9.6 | 0.846 |  | TFTC |  | 45.8±8.2 | 36.3±6.1 | 0.073 | **58.1±2.5** | **63.5±1.4** | **0.003** | |  | NA |  |
| CD11C+MHCII+ | 64.4±14.4 | 77.0±10.3 | 0.363 |  | TFTC |  | 78.8±4.1 | 60.0±22.3 | 0.101 | 64.2±3.7 | 69.6±6.7 | 0.151 | |  | NA |  |
| CD11C+MHClo | 79.3±2.6 | 78.5±6.7 | 0.810 |  | TFTC |  |  | TFTC |  | 81.7±8.0 | 90.9±4.2 | 0.054 | |  | NA |  |
| CD11C^neg^MHCII+ | 43.8±13.2 | 52.1±16.1 | 0.399 |  | TFTC |  | 43.0±11.9 | 41.4±9.9 | 0.820 | **38.1±4.3** | **46.4±6.0** | **0.034** | |  | NA |  |
| CD11C^neg^MHCII^neg^ | 36.2±10.5 | 36.6±9.1 | 0.591 |  | TFTC |  | 15.9±12.2 | 15.6±11.9 | 0.971 | 26.9±2.9 | 29.7±2.9 | 0.383 | |  | NA |  |
| CD11C+MHCII^neg^ | 71.9±6.6 | 71.8±8.6 | 0.975 |  | TFTC |  |  | TFTC |  | 87.5±4.5 | 90.0±7.4 | 0.543 | |  | NA |  |

All Macrophage populations are F480+. CD80 was used a surrogate for M1 macrophage phenotype. Average with SD. Students T-test. P< 0.05 highlighted. Jejunal, Ileal, and splenic specimen with n 10 (sham), 9 (SG). Data from these specimens represent combined results of two replicate experiments. Cecal samples with n 5 (sham), 5 (SG). TFTC (too few to count). CD80 was not available for the first panel iteration and thus hepatic populations were not surveilled for this marker (NA)

| **Supplemental Table 8: T cell subsets as % of CD45+ Leukocytes – DIO Sham versus SG** | | | | | | | | | | | | | | | | | | | | |  |  |  |
| --- | --- | --- | --- | --- | --- | --- | --- | --- | --- | --- | --- | --- | --- | --- | --- | --- | --- | --- | --- | --- | --- | --- | --- |
|  | **Jejunum** | | | | | **Ileum** | | | | | **Cecum** | | | | | **Spleen** | | | | | **Liver** | | |
|  | Sham | SG | | | p | Sham | SG | | | p | Sham | SG | | | p | Sham | | SG | | p | Sham | SG | P |
| CD3+ | 22.0±9.0 | | 18.5±9.6 | 0.417 | | 13.7±3.7 | | 17.8±8.6 | 0.366 | | 36.0±15.5 | | 34.1±22.0 | 0.882 | | **26.5±2.7** | **18.6±7.1** | | **0.009** | | 14.8±4.4 | 12.7±3.7 | 0.426 |
| CD3+103+ | 8.6±3.5 | | 6.9±4.8 | 0.403 | | 1.3±0.7 | | 2.8±2.8 | 0.294 | | 12.3±7.4 | | 8.4±8.1 | 0.469 | | 3.5±1.3 | 2.3±1.3 | | 0.060 | | 1.6±0.6 | 1.2±0.9 | 0.413 |
| CD3+CD69+ | 8.7±3.0 | | 6.9±3.8 | 0.270 | | 2.7±1.4 | | 3.9±2.5 | 0.367 | | 13.5±8.6 | | 9.9±9.3 | 0.564 | | 2.6±1.0 | 2.2±0.9 | | 0.312 | | 4.4±1.6 | 4.2±2.1 | 0.834 |
| CD4+ | 3.4±1.3 | | 2.6±1.5 | 0.211 | | 4.8±2.3 | | 8.1±3.0 | 0.098 | | 4.2±1.8 | | 3.1±0.8 | 0.303 | | **15.1±1.7** | **10.8±4.1** | | **0.013** | | 7.7±2.9 | 6.3±1.7 | 0.336 |
| CD4+CD103+ | 0.46±0.23 | | 0.31±0.12 | 0.080 | | 0.16±0.10 | | 0.44±0.42 | 0.189 | | 0.21±0.10 | | 0.21±0.16 | 0.993 | | 0.39±0.17 | 0.32±0.15 | | 0.373 | | 0.33±0.16 | 0.21±0.10 | 0.189 |
| CD4+FOXP3+ | 0.42±0.21 | | 0.33±0.14 | 0.260 | | 0.40±0.27 | | 1.3±1.7 | 0.287 | | 0.43±0.36 | | 0.53±0.27 | 0.684 | | 0.87±0.27 | 0.61±0.26 | | 0.116 | | 0.28±0.06 | 0.21±0.11 | 0.308 |
| CD4+GATA3+ | 0.55±0.54 | | 0.34±0.15 | 0.247 | | 0.21±0.11 | | 0.42±0.41 | 0.325 | | 0.33±0.35 | | 0.37±0.21 | 0.855 | | 1.4±1.6 | 1.2±1.3 | | 0.844 | | 2.3±1.1 | 1.7±0.7 | 0.333 |
| CD4+RORγt+ | 0.31±0.22 | | 0.21±0.10 | 0.202 | | 0.45±0.34 | | 1.3±1.8 | 0.316 | | 0.44±0.42 | | 0.51±0.28 | 0.782 | | 0.27±0.22 | 0.18±0.21 | | 0.365 | | 0.12±0.11 | 0.12±0.13 | 0.964 |
| CD4+Tbet+ | 0.15±0.11 | | 0.08±0.10 | 0.208 | | 0.27±0.17 | | 0.42±0.31 | 0.383 | | 0.34±0.37 | | 0.46±0.34 | 0.632 | | 0.28±0.27 | 0.12±0.19 | | 0.162 | | 0.01±0.00 | 0.01±0.01 | 0.883 |
| CD8+ | 13.4±7.6 | | 9.7±6.4 | 0.280 | | 2.6±1.3 | | 4.5±3.1 | 0.233 | | 11.3±5.7 | | 7.2±5.1 | 0.298 | | **10.0±1.2** | **6.3±2.7** | | **0.003** | | 4.1±0.9 | 3.3±1.3 | 0.272 |
| CD8+CD103+ | 6.9±3.1 | | 5.3±4.0 | 0.343 | | 0.73±0.53 | | 1.6±1.6 | 0.287 | | 6.0±3.7 | | 2.9±2.7 | 0.199 | | 3.31±1.3 | 2.2±1.4 | | 0.100 | | 1.0±0.3 | 0.76±0.63 | 0.453 |
| CD8+CD103+CD69+ | 4.4±2.1 | | 3.4±2.4 | 0.340 | | 0.28±0.21 | | 0.72±0.89 | 0.315 | | 3.7±2.6 | | 1.6±1.6 | 0.200 | | 0.30±0.33 | 0.28±0.25 | | 0.851 | | 0.74±0.04 | 0.59±0.26 | 0.308 |
| CD8+FoxP3+ | 0.36±0.24 | | 0.34±0.31 | 0.871 | | 0.19±0.10 | | 0.50±0.63 | 0.306 | | 0.25±0.20 | | 0.54±0.32 | 0.133 | | 0.08±0.04 | 0.07±0.05 | | 0.530 | | 0.07±0.04 | 0.07±0.06 | 0.915 |
| CD8+GATA3+ | 0.75±0.50 | | 0.44±0.31 | 0.126 | | 0.17±0.06 | | 0.48±0.64 | 0.298 | | 0.30±0.18 | | 0.38±0.22 | 0.538 | | 0.43±0.26 | 0.39±0.39 | | 0.821 | | 0.65±0.25 | 0.49±0.35 | 0.456 |
| CD8+Tbet+ | 1.3±1.3 | | 1.0±1.7 | 0.692 | | 0.26±0.15 | | 0.78±1.0 | 0.292 | | 0.90±0.59 | | 0.77±0.60 | 0.755 | | 0.48±0.45 | 0.17±0.28 | | 0.091 | | 0.01±0.01 | 0.01±0.00 | 0.652 |

Average with SD. Students T-test. P< 0.05 highlighted. Jejunal and splenic specimen with n 8 (sham), 10 (SG). Data from these specimens represents combined results of two replicate experiments. Liver specimen with n = 4 (sham), 6 (SG). Ileal and cecal samples with n 5 (sham), 4 (SG).

| **Supplemental Table 9: NKT, NK, IL cell subsets as % of CD45+ Leukocytes – DIO Sham versus SG** | | | | | | | | | | | | | | | | | | | | |  |  |  |
| --- | --- | --- | --- | --- | --- | --- | --- | --- | --- | --- | --- | --- | --- | --- | --- | --- | --- | --- | --- | --- | --- | --- | --- |
|  | **Jejunum** | | | | | **Ileum** | | | | | **Cecum** | | | | | **Spleen** | | | | | **Liver** | | |
|  | Sham | SG | | | p | Sham | SG | | | p | Sham | SG | | | p | Sham | | SG | | p | Sham | SG | P |
| CD3+NK1.1+ | 0.28±0.12 | | 0.22±0.24 | 0.503 | | 0.16±0.16 | | 0.50±0.58 | 0.238 | | 0.61±0.51 | | 0.45±0.48 | 0.640 | | 0.32±0.15 | 0.23±0.14 | | 0.176 | | 0.67±0.14 | 0.65±0.32 | 0.940 |
| CD3+NKp46+ | 0.70±0.53 | | 0.55±0.45 | 0.520 | | 0.34±0.30 | | 0.31±0.23 | 0.874 | | 1.0±1.0 | | 1.2±0.8 | 0.747 | | 0.47±0.24 | 0.35±0.30 | | 0.415 | | 0.54±0.14 | 0.33±0.34 | 0.271 |
| CD3+NK1.1+NKp46+ | 0.40±0.32 | | 0.24±0.23 | 0.211 | | 0.31±0.21 | | 0.86±0.92 | 0.231 | | 0.85±1.03 | | 1.0±1.0 | 0.782 | | 0.26±0.13 | 0.14±0.12 | | 0.052 | | 0.15±0.04 | 0.09±0.04 | 0.062 |
| CD3^neg^NK1.1+ | 0.74±0.34 | | 0.63±0.23 | 0.401 | | 0.40±0.14 | | 1.3±1.2 | 0.140 | | 0.75±0.31 | | 0.70±0.54 | 0.873 | | 0.56±0.21 | 0.71±0.30 | | 0.249 | | 3.5±1.5 | 2.5±1.6 | 0.363 |
| CD3^neg^NKp46+ | 1.6±1.9 | | 1.3±1.6 | 0.705 | | 1.1±0.9 | | 1.2±0.9 | 0.841 | | 2.9±3.2 | | 3.1±2.0 | 0.910 | | 1.0±0.5 | 1.8±1.8 | | 0.295 | | 2.8±1.2 | 1.9±0.4 | 0.119 |
| CD3^neg^NK1.1+NKp46+ | 0.48±0.38 | | 0.33±0.33 | 0.362 | | 0.30±0.18 | | 1.3±1.6 | 0.185 | | 2.0±1.8 | | 2.2±2.1 | 0.881 | | **2.4±0.6** | **1.7±0.4** | | **0.006** | | 2.9±1.3 | 2.1±0.5 | 0.200 |
| ILC1* (TBet+) | 0.56±0.60 | | 0.35±0.52 | 0.440 | | 0.28±0.18 | | 0.44±0.40 | 0.449 | | 0.50±0.34 | | 0.33±0.18 | 0.414 | | 0.14±0.15 | 0.01±0.10 | | 0.242 | | 0.03±0.01 | 0.02±0.01 | 0.208 |
| ILC2 (Gata3+) | 1.4±1.3 | | 1.2±0.5 | 0.608 | | 1.0±0.9 | | 0.84±0.64 | 0.767 | | 0.41±0.28 | | 0.41±0.22 | 1.00 | | 0.31±0.23 | 0.70±0.90 | | 0.257 | | 2.1±1.2 | 2.9±1.6 | 0.595 |
| ILC3 (RORγt+) | 1.0±1.0 | | 0.6±0.6 | 0.251 | | 1.1±0.7 | | 1.7±1.9 | 0.523 | | 0.44±0.32 | | 0.86±0.30 | 0.083 | | 0.10±0.10 | 0.19±0.19 | | 0.241 | | 0.71±0.79 | 0.78±0.76 | 0.885 |

Innate Lymphoid Cell. Derivative of a lineage negative population (CD45+CD3^neg^CD19^neg^CD11b^neg^CD11c^neg^NK1.1^neg^NKp46^neg^). Average with SD. Students T-test. P< 0.05 highlighted. Jejunal and splenic specimen with n 8 (sham), 10 (SG). Data from these specimens represents combined results of two replicate experiments. Liver specimen with n 4 (sham), 6 (SG). Ileal and cecal samples with n 5 (sham), 4 (SG).

| **Supplemental Table 10: CD19+11B**^n^**^eg^ B Cell subsets as % of CD45+ Leukocytes – DIO Sham versus SG** | | | | | | | | | | | | | | | | | | |  |  |  |
| --- | --- | --- | --- | --- | --- | --- | --- | --- | --- | --- | --- | --- | --- | --- | --- | --- | --- | --- | --- | --- | --- |
|  | **Jejunum** | | | **Ileum** | | | | | **Cecum** | | | | | **Spleen** | | | | | **Liver** | | |
|  | Sham | SG | p | Sham | SG | | | p | Sham | SG | | | p | Sham | | SG | | p | Sham | SG | P |
| Total | 17.3±14.3 | 11.4±6.1 | 0.247 | 42.2±19.1 | | 42.5±14.1 | 0.980 | | 29.3±16.7 | | 16.0±13.6 | 0.240 | | **59.2±1.6** | **45.2±13.1** | | **0.009** | | 32.6±6.3 | 25.7±9.3 | 0.234 |
| #Total CD80+ | 0.39±0.27 | 0.47±0.14 | 0.606 | 1.4±0.8 | | 1.6±0.6 | 0.797 | | 0.38±0.25 | | 0.21±0.01 | 0.218 | | 2.1±0.2 | 2.1±0.5 | | 0.949 | |  | NA |  |
| #Total IgM^neg^IgD+ | 3.3±4.3 | 3.1±1.3 | 0.904 | 9.5±4.2 | | 9.6±4.9 | 0.964 | | 5.3±2.9 | | 4.9±5.2 | 0.888 | | 9.5±3.4 | 9.1±4.0 | | 0.870 | |  | NA |  |
| #Total IgM+IgD^neg^ | 1.0±1.3 | 0.89±0.45 | 0.834 | 3.0±1.3 | | 3.2±0.7 | 0.771 | | 2.0±1.6 | | 1.7±1.1 | 0.709 | | 3.6±0.9 | 4.4±0.3 | | 0.154 | |  | NA |  |
| #Total IgM+IgD+ | 10.9±11.6 | 8.5±2.2 | 0.699 | 24.6±12.8 | | 24.2±8.3 | 0.955 | | 19.2±13.2 | | 6.1±4.8 | 0.104 | | **43.1±4.9** | **27.8±11.6** | | **0.030** | |  | NA |  |
| Total CD21+CD23^neg^ | 2.7±5.0 | 1.1±1.5 | 0.324 | 13.1±7.3 | | 9.7±5.2 | 0.452 | | 10.1±6.2 | | 4.8±4.6 | 0.198 | | 6.0±3.5 | 4.5±2.3 | | 0.305 | | **12.6±3.2** | **6.4±3.3** | **0.017** |
| Total CD21^neg^CD23+ | 0.68±0.69 | 1.0±0.7 | 0.312 | 1.0±0.6 | | 3.3±2.7 | 0.101 | | 0.59±0.44 | | 1.1±0.8 | 0.284 | | 1.6±0.6 | 2.3±1.5 | | 0.258 | | **0.64±0.20** | **1.3±0.5** | **0.038** |
| Total CD21+CD23+ | 10.1±8.0 | 6.1±3.4 | 0.174 | 19.5±12.2 | | 20.7±7.1 | 0.863 | | 14.9±14.8 | | 4.6±3.9 | 0.224 | | **45.9±3.4** | **30.8±10.9** | | **0.002** | | 14.5±8.6 | 14.1±6.0 | 0.933 |
| #CD21+CD23+IgM-IgD+ | 1.7±2.2 | 1.7±0.7 | 0.950 | 4.2±2.4 | | 4.6±2.3 | 0.784 | | 2.4±2.1 | | 1.5±1.5 | 0.483 | | 6.8±2.1 | 5.9±2.8 | | 0.599 | |  | NA |  |
| #CD21+CD23+IgM+IgD^neg^ | 0.23±0.23 | 0.23±0.13 | 1.000 | 0.49±0.28 | | 0.63±0.26 | 0.470 | | 0.31±0.14 | | 0.26±0.15 | 0.620 | | 1.2±0.3 | 1.2±0.1 | | 0.763 | |  | NA |  |
| #CD21+CD23+IgM+IgD+ | 7.1±7.1 | 5.9±1.4 | 0.754 | 13.9±9.3 | | 14.3±4.7 | 0.945 | | 11.6±12.3 | | 2.5±2.0 | 0.190 | | **35.2±5.7** | **21.0±9.6** | | **0.028** | |  | NA |  |

All B cell subsets from CD19+CD11b^neg^ parent. Average with SD. Students T-test. P< 0.05 highlighted. Jejunal and splenic specimen with n 8 (sham), 10 (SG) except as denoted by the symbol, “#”. Data from these specimens represents combined results of two replicate experiments. Liver specimen with n 4 (sham), 6 (SG) and for this experiment, IgD and CD80 was not in the staining panel (NA). Ileal and cecal samples with n 5 (sham), 4 (SG). #: data is strictly from the second panel iteration that included IgD and CD80 (n 5 for sham, 4 for SG).

| **Supplemental Table 11: CD19+11B+ B Cell subsets as % of CD45+ Leukocytes – DIO Sham versus SG** | | | | | | | | | | | | | | | | | | |  |  |  |
| --- | --- | --- | --- | --- | --- | --- | --- | --- | --- | --- | --- | --- | --- | --- | --- | --- | --- | --- | --- | --- | --- |
|  | **Jejunum** | | | **Ileum** | | | | | **Cecum** | | | | | **Spleen** | | | | | **Liver** | | |
|  | Sham | SG | p | Sham | SG | | | p | Sham | SG | | | p | Sham | | SG | | p | Sham | SG | P |
| Total | 0.38±0.49 | 0.41±0.49 | 0.899 | 0.02±0.02 | | 0.11±0.12 | 0.149 | | 0.07±0.12 | | 0.02±0.01 | 0.463 | | **1.5±0.4** | **3.7±2.3** | | **0.019** | | 0.81±0.45 | 0.73±0.29 | 0.748 |
| #Total CD80+ | 0.10±0.04 | 0.11±0.02 | 0.732 | 0.01±0.01 | | 0.05±0.06 | 0.179 | | 0.02±0.04 | | 0.01±0.01 | 0.400 | | 0.38±0.15 | 0.99±0.97 | | 0.201 | |  | NA |  |
| #Total IgM-IgD+ | 0.01±0.01 | 0.01±0.01 | 0.540 | **0.00±0.00** | | **0.01±0.00** | **0.007** | | 0.02±0.02 | | 0.00±0.00 | 0.421 | | 0.28±0.13 | 0.69±0.46 | | 0.096 | |  | NA |  |
| # Total IgM+IgD^neg^ | **0.02±0.02** | **0.04±0.02** | **0.049** | 0.00±0.00 | | 0.02±0.03 | 0.158 | | 0.00±0.01 | | 0.01±0.01 | 0.772 | | 0.13±0.04 | 0.49±0.43 | | 0.102 | |  | NA |  |
| # Total IgM+IgD+ | 0.02±0.02 | 0.03±0.02 | 0.661 | 0.01±0.01 | | 0.02±0.03 | 0.321 | | 0.03±0.07 | | 0.00±0.01 | 0.438 | | 0.61±0.24 | 1.1±0.9 | | 0.244 | |  | NA |  |
| Total CD21+CD23^neg^ | 0.05±0.05 | 0.07±0.10 | 0.735 | 0.00±0.00 | | 0.01±0.01 | 0.125 | | 0.01±0.03 | | 0.00±0.00 | 0.525 | | **0.26±0.07** | **0.56±0.33** | | **0.022** | | 0.23±0.13 | 0.17±0.04 | 0.342 |
| Total CD21^neg^CD23+ | 0.03±0.05 | 0.02±0.03 | 0.811 | 0.00±0.00 | | 0.00±0.00 | 0.214 | | 0 | | 0 | NA | | **0.06±0.03** | **0.26±0.20** | | **0.015** | | 0.03±0.02 | 0.03±0.02 | 0.736 |
| Total CD21+CD23+ | 0.11±0.20 | 0.12±0.17 | 0.938 | 0.00±0.00 | | 0.02±0.03 | 0.169 | | 0.03±0.07 | | 0.00±0.01 | 0.452 | | 0.76±0.34 | 1.7±1.4 | | 0.087 | | 0.36±0.26 | 0.20±0.09 | 0.209 |

All B cell subsets from CD19+CD11b^neg^ parent. Average with SD. Students T-test. P< 0.05 highlighted. Jejunal and splenic specimen with n 8 (sham), 10 (SG) except as denoted by the symbol, “#”. Data from these specimens represents combined results of two replicate experiments. Liver specimen with n 4 (sham), 6 (SG) and for this experiment, IgD and CD80 was not in the staining panel (NA). Ileal and cecal samples with n 5 (sham), 4 (SG). #: data is strictly from the second panel iteration that included IgD and CD80 (n 5 for sham, 4 for SG).

| **Supplemental Table 12: Myeloid Cell subsets as % of CD45+ Leukocytes – DIO Sham versus SG** | | | | | | | | | | | | | |  |  |  |
| --- | --- | --- | --- | --- | --- | --- | --- | --- | --- | --- | --- | --- | --- | --- | --- | --- |
|  | **Jejunum** | | | **Ileum** | | | **Cecum** | | | **Spleen** | | | | **Liver** | | |
|  | Sham | SG | p | Sham | SG | p | Sham | SG | p | Sham | SG | | p | Sham | SG | P |
| ***PMN***: Ly6G+ CD11b+ | 0.48±0.27 | 0.77±0.48 | 0.117 | 0.09±0.08 | 1.0±1.9 | 0.307 | **0.09±0.03** | **0.39±0.21** | **0.013** | **1.2±0.5** | **6.8±5.7** | **0.015** | | **0.71±0.23** | **0.34±0.14** | **0.020** |
| ***DC***: F480^neg^11C+MHCII+ | 1.1±0.9 | 0.63±0.43 | 0.196 | 0.14±0.11 | 0.28±0.28 | 0.349 | 0.24±0.13 | 0.32±0.28 | 0.611 | 0.27±0.17 | 0.41±0.43 | 0.404 | | 6.7±2.4 | 5.4±2.1 | 0.356 |
| ***MACS*** (Ly6G^neg^F480+) |  |  |  |  |  |  |  |  |  |  |  |  | |  |  |  |
| LY6C+ | 2.6±4.6 | 3.6±6.6 | 0.707 | 0.03±0.02 | 0.21±0.34 | 0.262 | 0.14±0.04 | 0.23±0.18 | 0.285 | 0.80±0.67 | 1.2±0.6 | 0.190 | | 2.6±0.6 | 2.3±0.9 | 0.690 |
| LYGC^neg^ | 11.3±8.4 | 11.5±8.3 | 0.957 | 0.34±0.30 | 0.92±1.15 | 0.308 | 0.30±0.14 | 0.29±0.17 | 0.917 | 0.66±0.29 | 0.97±0.45 | 0.103 | | 1.8±0.4 | 1.2±0.6 | 0.114 |
| CD11C+MHCII+ | 0.62±0.50 | 0.48±0.40 | 0.521 | 0.04±0.04 | 0.09±0.14 | 0.421 | 0.03±0.02 | 0.04±0.05 | 0.667 | 0.40±0.27 | 0.44±0.25 | 0.750 | | 0.50±0.18 | 0.45±0.23 | 0.710 |
| CD11C+MHClo | 4.6±5.2 | 3.6±3.4 | 0.630 | 0.05±0.06 | 0.14±0.18 | 0.316 | 0.03±0.02 | 0.01±0.01 | 0.225 | 0.24±0.15 | 0.38±0.28 | 0.218 | | 2.0±0.9 | 1.4±0.7 | 0.269 |
| CD11C-MHCII+ | 0.97±0.89 | 2.6±4.8 | 0.332 | 0.18±0.17 | 0.34±0.30 | 0.349 | 0.13±0.05 | 0.16±0.06 | 0.496 | 0.39±0.28 | 0.68±0.32 | 0.063 | | 1.5±0.7 | 1.4±0.5 | 0.196 |
| CD11C-MHCII- | 1.5±1.5 | 2.3±2.0 | 0.326 | 0.04±0.04 | 0.24±0.33 | 0.216 | 0.20±0.07 | 0.26±0.23 | 0.619 | 0.28±0.17 | 0.44±0.19 | 0.087 | | 0.51±0.17 | 0.64±0.37 | 0.558 |
| CD11C+MHCII- | 5.9±5.3 | 5.7±4.0 | 0.904 | 0.04±0.02 | 0.29±0.52 | 0.323 | 0.03±0.01 | 0.05±0.04 | 0.563 | 0.12±0.06 | 0.21±0.19 | 0.219 | | 2.3±0.8 | 1.7±1.1 | 0.374 |

All Macrophage populations are F480^+^. Average with SD. Students T-test. P< 0.05 highlighted. Jejunal and splenic specimen with n 8 (sham), 10 (SG). Data from these specimens represents combined results of two replicate experiments. Liver specimen with n 4 (sham), 6 (SG). Ileal and cecal samples with n 5 (sham), 4 (SG). Ly6C expression was measured from total macrophage parent.

| **Supplemental Table 13: CD80+ M1 Macrophage polarization as percent of Parent – DIO Sham versus SG** | | | | | | | | | | | | | |  |  |  |
| --- | --- | --- | --- | --- | --- | --- | --- | --- | --- | --- | --- | --- | --- | --- | --- | --- |
|  | **Jejunum** | | | **Ileum** | | | **Cecum** | | | **Spleen** | | | | **Liver** | | |
|  | Sham | SG | p | Sham | SG | p | Sham | SG | p | Sham | SG | | p | Sham | SG | P |
| MACS: Ly6G^neg^F480+ | 70.9±5.6 | 66.6±14.4 | 0.551 | 31.4±17.7 | 44.3±11.8 | 0.249 | 17.2±11.5 | 14.8±12.6 | 0.774 | 47.6±5.6 | 43.6±10.7 | 0.485 | |  | NA |  |
| CD11C+MHCII+ | 70.3±11.4 | 75.6±8.4 | 0.466 | 34.5±10.0 | 66.9±30.1 | 0.056 | 39.9±33.6 | 77.9±15.7 | 0.077 | 58.5±8.7 | 60.9±7.5 | 0.684 | |  | NA |  |
| CD11C+MHClo | 78.9±5.4 | 73.1±13.5 | 0.410 | 37.5±27.3 | 62.8±25.4 | 0.197 | 39.0±43.7 | 33.3±47.1 | 0.856 | 14.3±30.1 | 25.8±26.4 | 0.399 | |  | NA |  |
| CD11C^neg^MHCII+ | 41.8±7.8 | 43.4±11.3 | 0.809 | 23.1±11.1 | 45.7±17.6 | 0.051 | 21.1±16.9 | 17.8±22.4 | 0.808 | 22.4±4.9 | 21.0±8.2 | 0.768 | |  | NA |  |
| CD11C^neg^MHCII^neg^ | 39.7±4.9 | 44.6±14.8 | 0.502 | 17.1±8.0 | 23.7±6.5 | 0.219 | 3.7±2.4 | 3.7±5.8 | 0.993 | 14.8±3.3 | 15.4±7.5 | 0.878 | |  | NA |  |
| CD11C+MHCII^neg^ | 73.6±5.8 | 69.8±14.7 | 0.595 | 45.3±17.1 | 56.9±11.2 | 0.280 | **25.6±9.9** | **6.3±8.6** | **0.018** | 90.7±3.9 | 91.3±4.3 | 0.828 | |  | NA |  |

All Macrophage populations are F480+. CD80 was used a surrogate for M1 macrophage phenotype. Average with SD. Students T-test. P< 0.05 highlighted. Jejunal and splenic specimen with n 8 (sham), 10 (SG). Data from these specimens represents combined results of two replicate experiments. Ileal and cecal samples with n 5 (sham), 4 (SG). CD80 was not available for the first panel iteration and thus hepatic populations were not surveilled for this marker (NA).
